# The *Nefl*^E397K^ mouse model demonstrates muscle pathology and motor function deficits consistent with CMT2E

**DOI:** 10.1101/2025.02.02.636119

**Authors:** Dennis O. Pérez-López, Audrey A. Shively, F. Javier Llorente Torres, Roxanne Muchow, Zaid Abu-Salah, Mohammed T. Abu-Salah, Michael L. Garcia, Catherine L. Smith, Nicole L. Nichols, Monique A. Lorson, Christian L. Lorson

**Affiliations:** Department of Veterinary Pathobiology, College of Veterinary Medicine, University of Missouri, Columbia, MO 65211, USA; Bond Life Sciences Center, University of Missouri, Columbia, MO 65211, USA; Department of Biological Sciences, College of Arts and Science, University of Missouri, Columbia, MO 65211, USA; Department of Biomedical Sciences, College of Veterinary Medicine, University of Missouri, Columbia, MO 65211, USA; Department of Medical Pharmacology and Physiology, School of Medicine, University of Missouri, Columbia, MO, USA; Dalton Cardiovascular Research Center, University of Missouri, Columbia, MO, USA

**Author notes:** Denotes co-corresponding authors: Christian L. Lorson and Monique A. Lorson, University of Missouri, Department of Veterinary Pathobiology, Bond Life Sciences Center, 1201 Rollins Street, Columbia, MO 65211. phone number: 1+(.

## Abstract

Charcot-Marie-Tooth (CMT) disease affects approximately 1 in 2,500 people and represents a heterogeneous group of inherited peripheral neuropathies characterized by progressive motor and sensory dysfunction. CMT type 2E is a result of mutations in the neurofilament light (*NEFL*) gene with predominantly autosomal dominant inheritance, often presenting with a progressive neuropathy with distal muscle weakness, sensory loss, gait disturbances, foot deformities, reduced nerve conduction velocity (NCV) without demyelination and typically reduced compound muscle action potential (CMAP) amplitude values. Several *Nefl* mouse models exist that either alter the mouse *Nefl* gene or overexpress a mutated human *NEFL* transgene, each recapitulating various aspects of CMT2E disease. We generated the orthologous *NEFL^E396K^* mutation in the mouse C57BL/6 background, *Nefl^E397K^*. In a separate report, we extensively characterized the electrophysiology deficits and axon pathology in *Nefl^E397K^* mice. In this manuscript, we report our characterization of *Nefl^E397K^* motor function deficits, muscle pathology and changes in breathing *Nefl^+/E397K^* and *Nefl^E397K/E397K^* mice demonstrated progressive motor coordination deficits and muscle weakness through the twelve months of age analyzed, consistent with our electrophysiology findings. Additionally, *Nefl^+/E397K^* and *Nefl^E397K/E397K^* mice showed alterations in muscle fiber area, diameter and composition as disease developed. Lastly, *Nefl* mutant mice showed increased number of apneas under normoxia conditions and increased erratic breathing as well as tidal volume under respiratory challenge conditions. *Nefl^E397K/E397K^* mice phenotypes and pathology were consistently more severe than *Nefl^+/E397K^* mice. Collectively, these novel CMT2E models present with a clinically relevant phenotype and make it an ideal model for the evaluation of therapeutics.

## Introduction

Charcot-Marie-Tooth (CMT) diseases are slowly progressing neuropathies with an incidence of 1 in 2,500, affecting around 3 million people worldwide. CMT2 is a slow, but progressive disorder associated with axonal dysfunction and the deterioration of axonal connections and communication with muscles leading to motor function impairment, balance deficits, muscle atrophy, and denervation.

Mutations in the neurofilament light gene (*NEFL*) are associated with axonal CMT2E and represent ∼1% of all CMT cases. Clinical symptoms are variable in the age of onset and the severity but largely include progressive distal muscle weakness, sensory loss, foot deformities, gait disturbances and altered nerve conduction velocity (NCV) and compound muscle action potential (CMAP). The majority of *NEFL* mutations are inherited in a dominant manner; recessive mutations are rare (1-3). There are over thirty different *NEFL* mutations that are predominantly found within the functional domains of the NF-L protein (head, rod and tail) and are predicted to disrupt neurofilament assembly. The *NEFL^E396K^* mutation is one of the most identified patient mutations (4-9).

*NEFL* encodes the neurofilament light (NF-L) protein. NF-L is one of the core intermediate filament components including neurofilament medium (NF-M) and neurofilament heavy (NF-H). Peripherin is also a component of the neurofilament core within the peripheral nervous system (PNS) while alpha-internexin is a core component of the central nervous system (CNS). These core components assemble into an intermediate filament that serves multiple functions including structural stability, axonal growth, nerve conduction and transport. Mutations within these core proteins disrupt neurofilament assembly and often lead to intermediate filament aggregation (6, 10-14).

There are several mouse models for *NEFL* (*Nefl^N98S^*, *Nefl^P8R^*, *Nefl^L394P^*, h*NEFL^E396K^* and h*NEFL^P22S^*) (15-21). *Nefl^N98S^* and *Nefl*^P8R^ models have the orthologous mutation generated within the mouse genome while the *Nefl^L394P^* mouse has mutant mouse *Nefl* driven by the murine sarcoma virus promoter. h*NEFL^E396K^* and h*NEFL^P22S^* models overexpress the human *E396K* or *P22S* transgenes, respectively, in the context of wild type mouse *Nefl* expression. Of these models, the *Nefl^N98S^* model demonstrates the most severe phenotypes based on age of onset (6+ weeks) and disease pathology. At six weeks, mutant mice develop progressive hindlimb weakness, altered balance and gait disturbances. A reduction in CMAP and NCV was observed as well as smaller myelinated axons, thicker myelin sheaths, fewer neurofilaments in axons, and neurofilament aggregates (15, 16). h*NEFL^E396K^* mice develop hindlimb weakness and motor coordination deficits at ∼4 months of age. h*NEFL^E396K^* mice demonstrate neurofilament accumulation, reduced axon diameters, reduced nerve conduction velocities (NCV) and muscle fiber alterations without neuromuscular junction (NMJ) denervation (17, 18). At 6 months of age, h*NEFL^P22S^* mice developed hindlimb weakness, gait and coordination disturbances and muscle pathology with no axonal loss; however, unlike h*NEFL^E396K^* mice, h*NEFL^P22S^* mice also demonstrated NMJ denervation (19, 21). *Nefl^L394P^* mice develop hindlimb weakness and abnormal gait with severe and progressive disease with death largely within 28 days. Axonal degeneration, neurofilament accumulation and muscle fiber atrophy were observed (20).

We generated the *NEFL^E396K^* mutation within the context of the mouse genome, C56BL/6-*Nefl*^E397K^, to examine CMT2E disease progression and to develop a model to readily evaluate therapeutic efficacy. While *Nefl^E397K^* is inherited in a dominant manner, we examined *Nefl^+/E397K^* and *Nefl^E397K/E397K^* mice to understand whether disease progression and severity were altered within the two genetic contexts. A companion manuscript details the extensive characterization of *Nefl^+/E397K^* and *Nefl^E397K/E397K^* electrophysiology and axon pathology and demonstrates that these mouse models demonstrate quantitative differences as early as three weeks that are consistent with CMT2E. In this manuscript, we analyze *Nefl*^+/E397K^ and *Nefl*^E397K/E397K^ motor function, muscle pathology and respiration and report progressive motor function and coordination deficits that are consistent with our previously reported electrophysiology findings. *Nefl^+/E397K^* and *Nefl^E397K/E397K^* mice showed alterations in muscle fibers as disease developed, as well as differences in apneas, erratic breathing and tidal volume. Together, these manuscripts provide an extensive characterization of the *Nefl-E397K* mouse models and demonstrate their utility in evaluating therapeutic efficacy.

## Results

### *Nefl* mutants demonstrate motor coordination, gait and balance deficits

CMT2E patients present with motor coordination, gait, balance deficits and progressive muscle weakness. We evaluated motor function and coordination using hindlimb splay (HLS), time-to-right (TTR), grip strength, rotarod, dowel rod and CatWalk assessments. HLS was scored 0-3: a score of 3 indicated fully extended hindlimbs; this was a routine score for wild type mice. Mice scored 2 if the hindlimbs extended to a hip-width distance and scored 1 if hindlimbs were positioned near midline. Mice scored 0 when hindlimbs were completely retracted or were clasped together. During the 360-day assessment period, *Nefl^+/E397K^* and *Nefl^E397K/E397K^* mice did not show significant differences from wild type mice in hindlimb splay (Fig. 1A). TTR was conducted to analyze early core strength. *Nefl^+/E397K^* and *Nefl^E397K/E397K^* mice did not show differences in TTR compared to wild type littermates during the thirty days of assessment (Fig. 1B). *Nefl^+/E397K^* and *Nefl^E397K/E397K^* mice muscle strength and balance were further analyzed on a rotarod apparatus. *Nefl^+/E397K^* and *Nefl^E397K/E397K^* mice showed significant differences compared to the wild type cohort in time spent walking on the rotarod throughout the 180 day assessment period (WT mean average=111.4 seconds, *Nefl^+/E397K^* mean average=79.7 seconds (*P*=0.0001), *Nefl^E397K/E397K^* mean average=81.9 seconds (*P*=0.0002)) (Fig. 1C). To assess muscle strength between wild type and *Nefl* mutant mice, hindlimb and all-limb grip strength were measured. There were significant differences observed in all-limb grip strength between *Nefl^+/E397K^*, *Nefl^E397K/E397K^* and wild type mice during the assessment period (WT mean average=151.3 grams, *Nefl^+/E397K^* mean average=143.0 grams (*P*=0.0013), *Nefl^E397K/E397K^* mean average=135.4 grams (*P*<0.0001)) (Fig. 1D). In contrast, there were no significant difference in hindlimb grip strength for either *Nefl^+/E397K^* or *Nefl^E397K/E397K^* mice when compared to wild type littermates (Fig. 1E).

**Figure 1.**
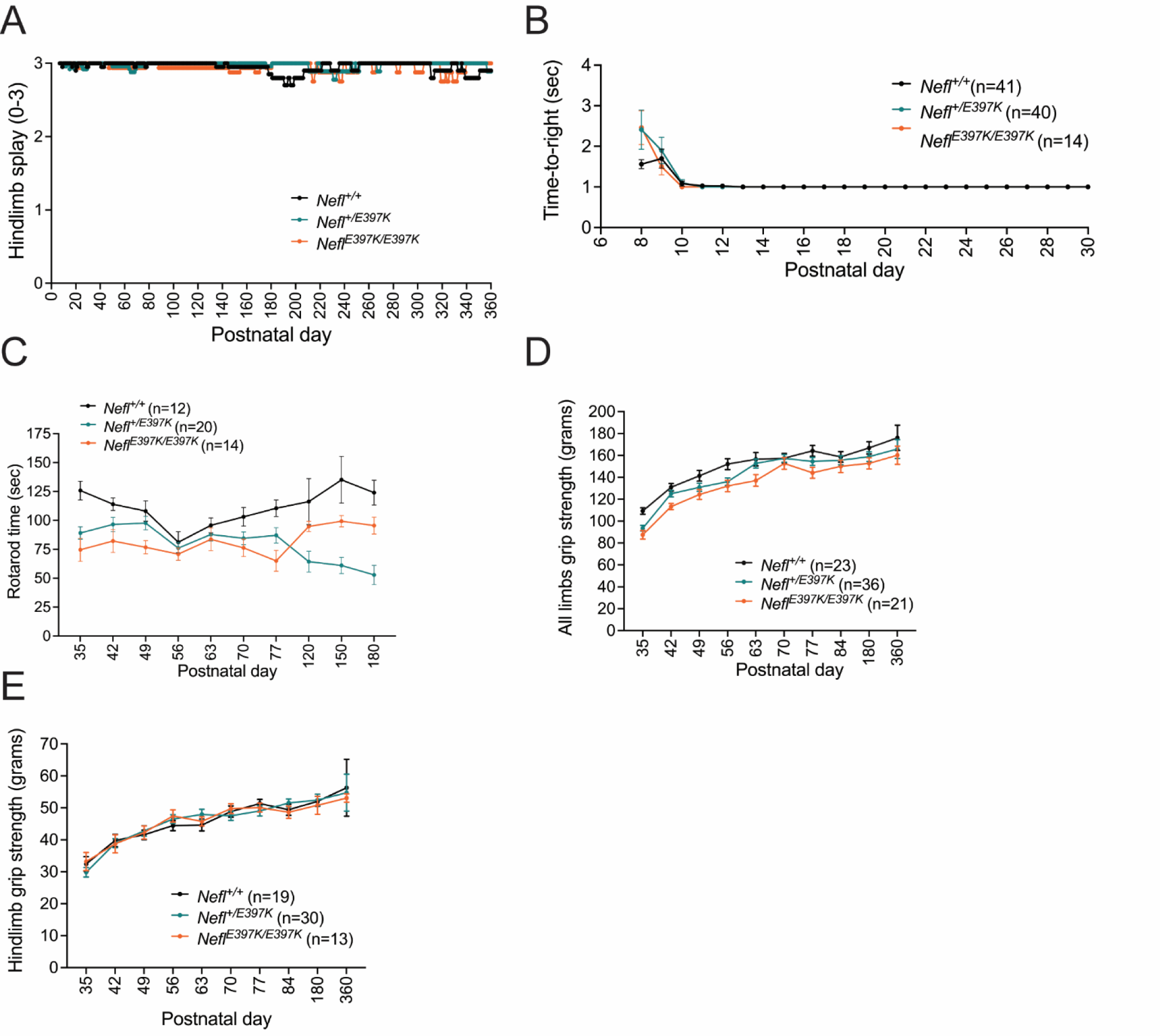
*Nefl* mutants show reduced motor function and grip strength. (A) Hindlimb splay score (mean ± SEM) from P7-P360 for *Nefl^+/+^* (black, mean=3.0, N=68), *Nefl^+/E397K^* (teal, mean=3.0, N=85), and *Nefl^E397K/E397K^* (orange, mean=2.9, N=35). There were not significant differences between *Nefl* mutant and wild type cohorts. (B) Time-to-right from P8-P30 measured in seconds (mean ± SEM) for *Nefl^+/+^* (black, mean=1.1, N=41), *Nefl^+/E397K^* (teal, mean=1.1, N=40), and *Nefl^E397K/E397K^* (orange, mean=1.1, N=14). There were not significant differences between *Nefl* mutant and wild type cohorts. (C) Rotarod measured in seconds (mean ± SEM) for *Nefl^+/+^* (black, mean=111.4, N=12), *Nefl^+/E397K^* (teal, mean=79.7, *P*<0.0001, N=20), and *Nefl^E397K/E397K^* (orange, mean=81.9, *P*=0.0002, N=14). (D) All limbs grip strength measured in grams (mean ± SEM) for *Nefl^+/+^* (black, mean=151.3, N=23), *Nefl^+/E397K^* (teal, mean=143.0, *P*=0.0013, N=36), and *Nefl^E397K/E397K^* (orange, mean=135.4, *P*<0.0001, N=21). (E) Hindlimb grip strength measured in grams (mean ± SEM) for *Nefl^+/+^* (black, mean=46.1, N=19), *Nefl^+/E397K^* (teal, mean=46.1, N=30), and *Nefl^E397K/E397K^* (orange, mean=46.0, N=13). Two-way ANOVA with Dunnett multiple comparison test was used for statistical analyses in A and B. Ordinary one-way ANOVA with Dunnett multiple comparison test was used to determine significance in C. Repeated measured one-way ANOVA with Dunnett multiple comparison test was used to determine significance in D and E. N = number of mice evaluated.

A longitudinal assessment of gait, coordination, balance, and motor function were performed using a dowel rod test for *Nefl^+/E397K^* and *Nefl^E397K/E397K^* mice. Consistent with progressive motor coordination deficits, there was a significant difference between wild type and *Nefl^E397K/E397K^* mice in the time to traverse the dowel rod (WT mean=11.3 seconds, *Nefl^+/E397K^* mean=12.1 seconds (NS), *Nefl^E397K/E397K^* mean=14 seconds (*P*=0.0343)). At P360, the latest time point measured, there were significant differences between wild type (7.2 seconds) and *Nefl^+/E397K^* and *Nefl^E397K/E397K^* mice (14.8 seconds, *P*=0.0159; 14.0 seconds, *P*=0.0097, respectively) (Fig. 2A), suggesting a progressive loss of motor function. There were no significant differences between the three cohorts in the number of steps traversed across the dowel rod but there were significant differences in gait (steps/second) as disease progressed (Fig. 2B, C). At P360, gait was significantly shortened in *Nefl* mutants (WT mean=2.44 steps/second, *Nefl^+/E397K^* mean=0.98 steps/second (*P*=0.0215), *Nefl^E397K/E397K^* mean=1.1 steps/second (*P*=0.0195)) (Fig. 2C). Balance was assessed by scoring the number of tail grabs, foot slips and the tail position. Tail position was scored 1-3 where 1 indicated the tail was used to balance and support, a score of 2 indicated the tail position was parallel to the dowel, and score of 3 indicated the tail position was raised vertical. There was not a significant difference in the average tail position between wild type and *Nefl^+/E397K^* mice (WT mean=1.7, *Nefl^+/E397K^* mean=1.6, NS) but there was a difference between wild type and *Nefl^E397K/E397K^* mice (*Nefl^E397K/E397K^* mean=1.3, *P*=0.0299). Disease progression influenced tail position between wild type, *Nefl^+/E397K^* and *Nefl^E397K/E397K^* mice suggesting increased motor coordination deficits (P360 WT=1.8, *Nefl^+/E397K^*=1.0, NS, *Nefl^E397K/E397K^*=0.8, *P*=0.0048) (Fig. 2D). The number of tail grabs (P360 WT=7.3, *Nefl^+/E397K^*=15.3, *P*=0.0322, *Nefl^E397K/E397K^*=12.4, NS) and the number of foot slips (P360 WT=4.7, *Nefl^+/E397K^*=18.5, NS, *Nefl^E397K/E397K^*=10.3, NS) increased in *Nefl* mutants as disease progressed supporting that motor coordination deficits were increased (Fig. 2E-F). *Nefl^+/E397K^* and *Nefl^E397K/E397K^* mice were observed to drag their bodies across the dowel, place their paws on each side of the dowel and their tails were more frequently wound around the dowel rod for movement. In contrast, wild type littermates typically traversed the dowel with their tail raised vertical and their paws placed on the dowel rod not around the dowel rod.

**Figure 2.**
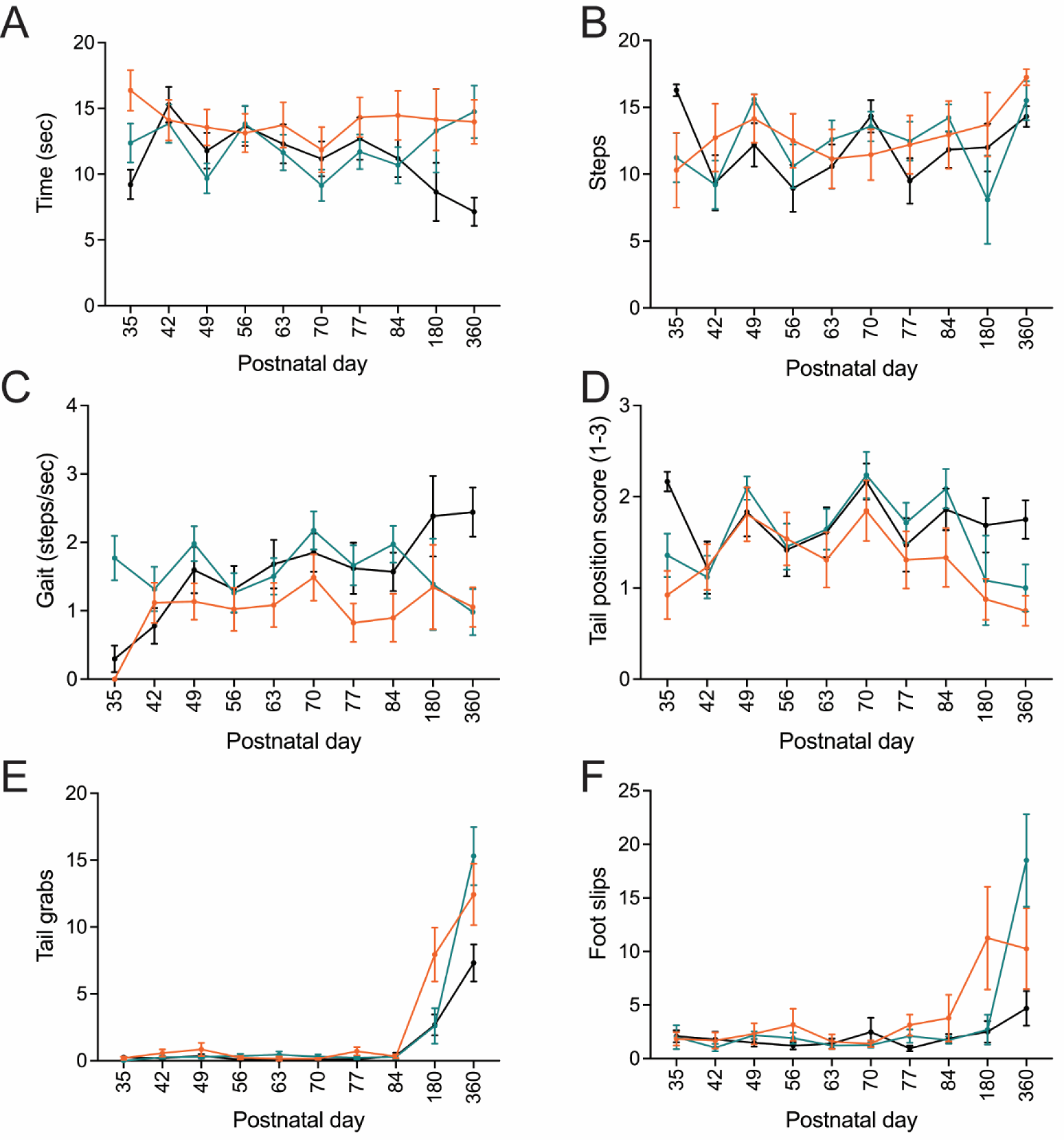
*Nefl* mutant mice show motor coordination deficits. Longitudinal dowel rod assessment was performed on *Nefl^+/+^* (black, N=23), *Nefl^+/E397K^* (teal, N=36), and *Nefl^E397K/E397K^* (orange, N=16) mice. (A) Average time to traverse the dowel (seconds) for *Nefl^+/+^* (P360 mean=7.2), *Nefl^+/E397K^* (mean=14.8, *P*=0.0159), and *Nefl^E397K/E397K^* (mean=14.0, *P*=0.0097). (B) Average number of steps to traverse the dowel for *Nefl^+/+^* (mean=14.3), *Nefl^+/E397K^* (mean=16.9, *P*=0.0551), and *Nefl^E397K/E397K^* (mean=17.3, *P*=0.0216). (C) Average gait (steps/seconds) for *Nefl^+/+^* (mean=2.4), *Nefl^+/E397K^* (mean=1.0, *P*=0.0215), and *Nefl^E397K/E397K^* (mean=1.1, *P*=0.0195). (D) Balance assessment by tail position score for *Nefl^+/+^* (mean=1.8), *Nefl^+/E397K^* (mean=1.0, NS), and *Nefl^E397K/E397K^* (mean=0.8, *P*=0.0048). (E) Average number of tail grabs for *Nefl^+/+^* (mean=7.3), *Nefl^+/E397K^* (mean=15.3, *P*=0.0332), and *Nefl^E397K/E397K^* (mean=12.4, NS). (F) Average number of foot slips for *Nefl^+/+^* (mean=4.7), *Nefl^+/E397K^* (mean=18.5, NS), and *Nefl^E397K/E397K^* (mean=10.3, NS). Two-way ANOVA with Dunnett multiple comparisons test was used to determine significance. N=number of mice evaluated, NS=not significant.

Numerous gait and motor function deficits were also measured using an automated gait analysis CatWalk system. Catwalk parameters that were significantly different for CMT2E model mice were speed, base of support, maximal contact area, print width and print position with *Nefl^E397K/E397K^* mice showing the largest differences from wild type mice (Table 1, Supplementary Fig. S1). Speed was recorded as the time to traverse the platform base. The base of support recorded the average width between the front and hind paws of an animal while walking. The maximal contact area was recorded as the time during a stance that the footprint was the largest in seconds. The print width measured the width of the complete footprint while the print position measured the position of the hind paw relative to the previous position of the fore paw. For speed, base of support, maximal contact area of the hind paws, print width and print position *Nefl^E397K/E397K^* mice demonstrated significant differences from wild type mice with *Nefl^+/E397K^* showing some differences (Table 1, Supplementary Fig. S1). Speed was routinely slower throughout the 360 days of assessment for *Nefl^E397K/E397K^* mice. For *Nefl^E397K/E397K^* mice, base of support and maximal contact area the hind paws showed the most variance from wild type mice when compared to the front paws (Table 1, Supplementary Fig. S1). As disease progressed, left and right print positions demonstrated more variance in *Nefl^E397K/E397K^* mice (Supplementary Fig. S1).

**Table 1.**

| **Table 1** | *Nefl*<sup>+/+</sup> | *Nefl*<sup>+/E397K</sup> | *Nefl*<sup>E397KE397K</sup> |
| --- | --- | --- | --- |
| Speed (time in seconds) | 17.05 | 19.46 | 14.27 |
| Base of support front paws | 1.38 | 1.48 | 1.66 |
| Base of support hind paws | 2.91 | 2.81 | 3.26 |
| Max contact area front L | 0.57 | 0.61 | 0.65 |
| Max contact area front R | 0.69 | 0.68 | 0.44 |
| Max contact area hind L | 0.60 | 0.73 | 0.53 |
| Max contact area hind R | 0.69 | 0.68 | 0.44 |
| Print width hind R | 0.95 | 0.94 | 0.58 |
| Print width hind L | 0.96 | 1.02 | 0.72 |
| Print width front L | 0.83 | 0.90 | 0.95 |
| Print width front R | 0.96 | 0.91 | 0.98 |
| Print position right side | 1.47 | 1.25 | 3.12 |
| Print position left side | 1.70 | 1.13 | 2.90 |

Catwalk analyses of wild type, *Nefl*<sup>+/E397K</sup>, and *Nefl*<sup>E397KE397K</sup> mice. The values reported are the average at P360.

### *Nefl* mutants show forelimb and hindlimb differences in muscle fibers

To evaluate muscle pathology in *Nefl^+/E397K^* and *Nefl^E397K/E397K^* mice, forelimb muscles (biceps brachii and triceps brachii) and hindlimb muscles (tibialis anterior and gastrocnemius) were analyzed at twelve weeks when differences in motor function were becoming prevalent (Fig. 3, Supplementary Fig. S2). Muscle fiber area (WT mean=1957μm^2^, *Nefl^+/E397K^* mean=1549μm^2^, *P*<0.0001, *Nefl^E397K/E397K^* mean=1655μm^2^, *P*<0.0001) and minimal Feret diameter (WT mean=45.1μm, *Nefl^+/E397K^* mean=39.4μm, *P*<0.0001, *Nefl^E397K/E397K^* mean=41.1μm, *P*<0.0001) were significantly reduced in *Nefl* mutants at 12 weeks in the biceps brachii (Fig. 3A,B). Interestingly, in *Nefl* mutants, triceps brachii muscle fiber area was similar to or larger than muscle fibers of wild type mice (WT mean=1769μm^2^, *Nefl^+/E397K^* mean=1929μm^2^, *P*=0.0012, *Nefl^E397K/E397K^* mean=1731μm^2^, NS) (Fig. 3C). There were no significant differences in triceps brachii minimal Feret diameter between wild type and *Nefl* mutant mice (Fig. 3D). Similar to the biceps brachii, the gastrocnemius showed significant reduction in muscle fiber area (WT mean=2826μm^2^, *Nefl^+/E397K^* mean=2304μm^2^, *P*<0.0001, *Nefl^E397K/E397K^* mean=2228μm^2^, *P*<0.0001) and minimal Feret diameter at twelve weeks (WT mean=53.2μm, *Nefl^+/E397K^* mean=48.0μm, *P*<0.0001, *Nefl^E397K/E397K^* mean=47.3μm, *P*<0.0001) (Fig. 3E,F). In the tibialis anterior, *Nefl^E397K/E397K^* muscle fiber area and minimal Feret diameter were slightly increased when compared to wild type mice at twelve weeks (Fig. 3G, H). In contrast, muscle fiber area and diameter were decreased in *Nefl^+/E397K^* mice when compared to wild type mice (WT mean area=2672μm^2^, *Nefl^+/E397K^* mean area=2478μm^2^, *P*=0.0038) (WT mean diameter=50.7μm, *Nefl^+/E397K^* mean=48.7μm, *P*=0.0059) (Fig. 3G, H). When the distribution of muscle fiber area was analyzed between wild type and *Nefl* mutant mice at twelve weeks, there were significant differences most notably in the biceps brachii with also significant differences noted in the gastrocnemius and tibialis anterior muscle fiber area (Supplementary Fig. S3).

**Figure 3.**
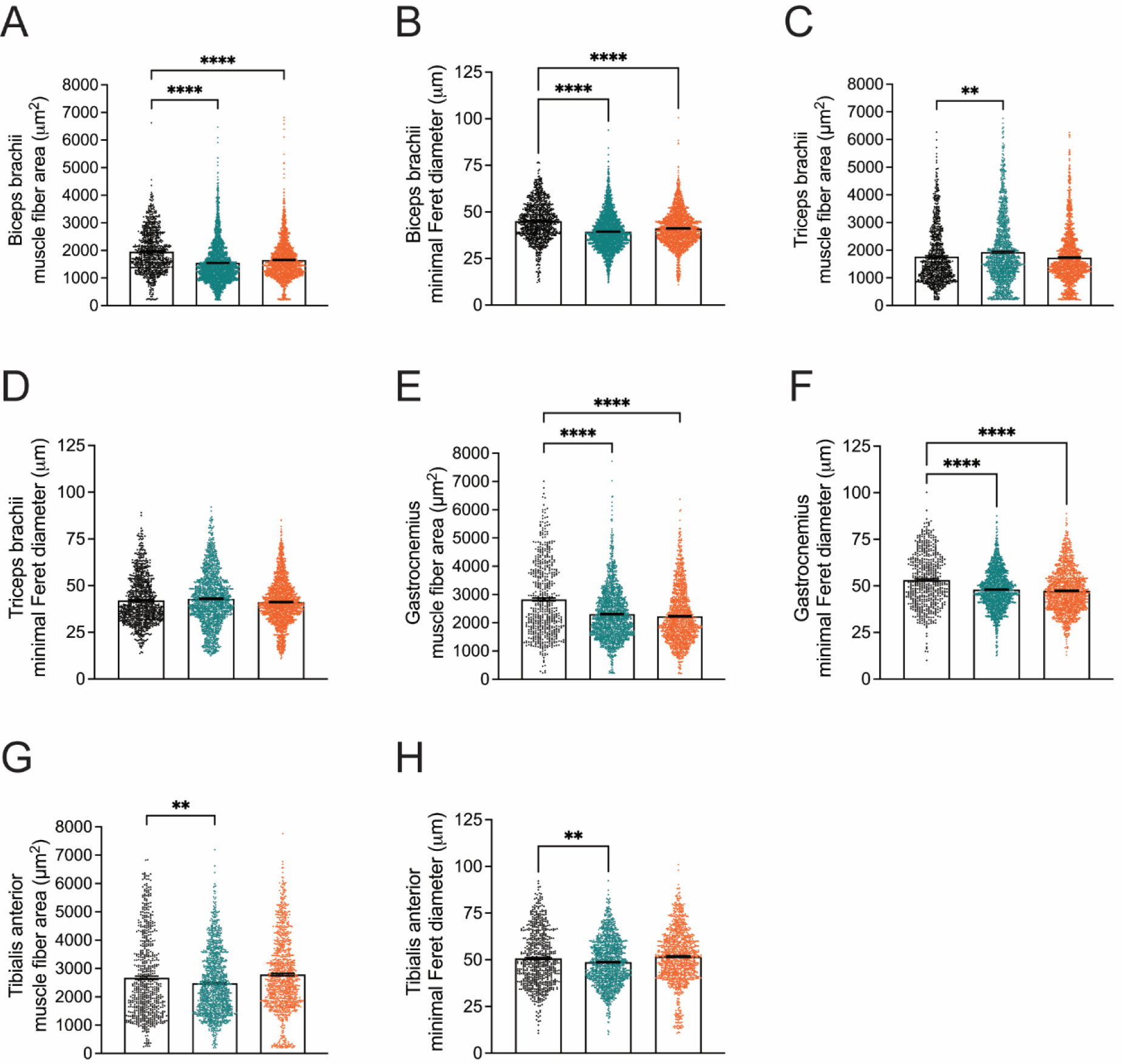
Muscle pathology in CMT2E mice show muscle defects at twelve weeks. Quantification of biceps brachii, triceps brachii, gastrocnemius, and tibialis anterior muscle fiber area for *Nefl^+/+^* (black), *Nefl^+/E397K^* (teal), and *Nefl^E397K/E397K^* (orange). (A) Biceps brachii area of *Nefl^+/+^*=1957μm^2^ (900 fibers, N=3), *Nefl^+/E397^*=1549μm^2^ (2335 fibers, *P*<0.0001, N=6), and *Nefl^E397K/E397K^*=1655μm^2^ (1695 fibers, *P*<0.0001, N=5). (B) Biceps brachii minimal Feret diameter of *Nefl^+/E397K^*=45.1μm (900 fibers, N=3), *Nefl^+/E397K^*=39.4μm (2335 fibers, *P*<0.0001, N=6), and *Nefl^E397K/E397K^*=41.1μm (1695 fibers, *P*<0.0001, N=5). (C) Triceps brachii area of *Nefl^+/+^*=1769μm^2^ (898 fibers, N=3), *Nefl^+/E397K^*=1929μm^2^ (1349 fibers, *P*=0.0012, N=5), and *Nefl^E397K/E397K^*=1731μm^2^ (1588 fibers, NS, N=5). (D) Triceps brachii minimal Feret diameter of *Nefl^+/+^*=42.0μm (898 fibers, N=3), *Nefl^+/E397K^*=42.9μm (1349 fibers, NS, N=5), and *Nefl^E397K/E397K^*=41.2μm (1588 fibers, NS, N=5). (E) Gastrocnemius area of *Nefl^+/+^*=2826 μm^2^ (565 fibers, N=3), *Nefl^+/E397K^*=2304μm^2^ (1496 fibers, *P*<0.0001, N=6), and *Nefl^E397K/E397K^*=2228μm^2^ (1284 fibers, *P*<0.0001, N=5). (F) Gastrocnemius minimal Feret diameter of *Nefl^+/+^*=53.2μm (565 fibers, N=3), *Nefl^+/E397K^*=48.0μm (1496 fibers, *P*<0.0001, N=6), and *Nefl^E397K/E397K^*=47.3μm (1284 fibers, *P*<0.0001, N=5). (G) Tibialis anterior area of *Nefl^+/+^*=2672μm^2^ (655 fibers, N=3), *Nefl^+/E397K^*=2478 μm^2^ (1179 fibers, *P*=0.0038, N=5), and *Nefl^E397K/E397K^*=2786μm^2^ (1036 fibers, NS, N=5). (H) Tibialis anterior minimal Feret diameter of *Nefl^+/+^*=50.7μm (655 fibers, N=3), *Nefl^+/E397K^*=48.7μm (1179 fibers, *P*=0.0059, N=5), and *Nefl^E397K/E397K^*=51.8μm (1036 fibers, NS, N=5). Ordinary one-way ANOVA with Dunnett multiple comparisons test was used to determine significance. N=number of mice evaluated, NS=not significant.

At twelve months of age, *Nefl* mutant mice demonstrated deficits in motor coordination, balance and gait, especially as it pertained to the hindlimb assessments; therefore, the same forelimb (bicep brachii and triceps brachii) and hindlimb muscles (tibialis anterior and gastrocnemius) were analyzed at twelve months (Fig. 4, Supplementary Fig. S2). Interestingly, while the biceps brachii demonstrated reduced muscle fiber area in both *Nefl* mutants at twelve weeks, muscle fiber area was similar or increased in the *Nefl* mutants at twelve months (WT mean=2032μm^2^, *Nefl^+/E397K^* mean=2276μm^2^, *P*<0.0001, *Nefl^E397K/E397K^* mean=2069μm^2^, NS) (Fig. 4A). Similarly, the biceps brachii minimal Feret diameter was increased at twelve months (WT mean=43.8μm, *Nefl^+/E397K^* mean=44.2μm, NS, *Nefl^E397K/E397K^* mean=45.8μm, *P*=0.0040) (Fig. 4B). In the triceps brachii (12M), there was a reduction in muscle fiber area and minimal Feret diameter in *Nefl^+/E397K^* mice when compared to wild type mice while *Nefl^E397K/E397K^* mice showed no significant differences (Fig. 4C, D). At twelve weeks, gastrocnemius muscle fibers were significantly reduced in area and diameter in *Nefl* mutants (Fig. 3E, F). In contrast, at twelve months, *Nefl^+/E397K^* and *Nefl^E397K/E397K^* mice showed significantly increased gastrocnemius muscle fiber area (WT mean=1984μm^2^, *Nefl^+/E397K^* mean=2349μm^2^, *P*<0.0001, *Nefl^E397K/E397K^* mean=2350μm^2^, *P*<0.0001) and diameter (WT mean=43.5μm, *Nefl^+/E397K^* mean=47.1μm, *P*<0.0001, *Nefl^E397K/E397K^* mean=46.3μm, *P*=0.0013) when compared to wild type mice (Fig. 4E, F). In contrast to twelve weeks, the tibialis anterior muscle fiber area and diameter in *Nefl^+/E397K^* mice was consistent with wild type mice at twelve months (WT mean area=2716μm^2^, *Nefl^+/E397K^* mean area=2671μm^2^, NS) (WT mean diameter=50.6μm, *Nefl^+/E397K^* mean diameter=49.9μm, NS) (Fig. 4G, H). At twelve months, *Nefl^E397K/E397K^* mice further increased muscle fiber area and diameter compared to wild type mice (WT mean area=2716μm^2^, *Nefl^E397K/E397K^* mean=3035μm^2^, *P*<0.0001) (WT mean diameter=50.6μm, *Nefl^E397K/E397K^* mean diameter=52.4μm, *P*=0.0257) (Fig. 4G. H). When the distribution of muscle fiber area was analyzed between wild type and *Nefl* mutant mice at twelve months, there were only a few significant differences noted in the biceps brachii (Supplementary Fig. S4).

**Figure 4.**
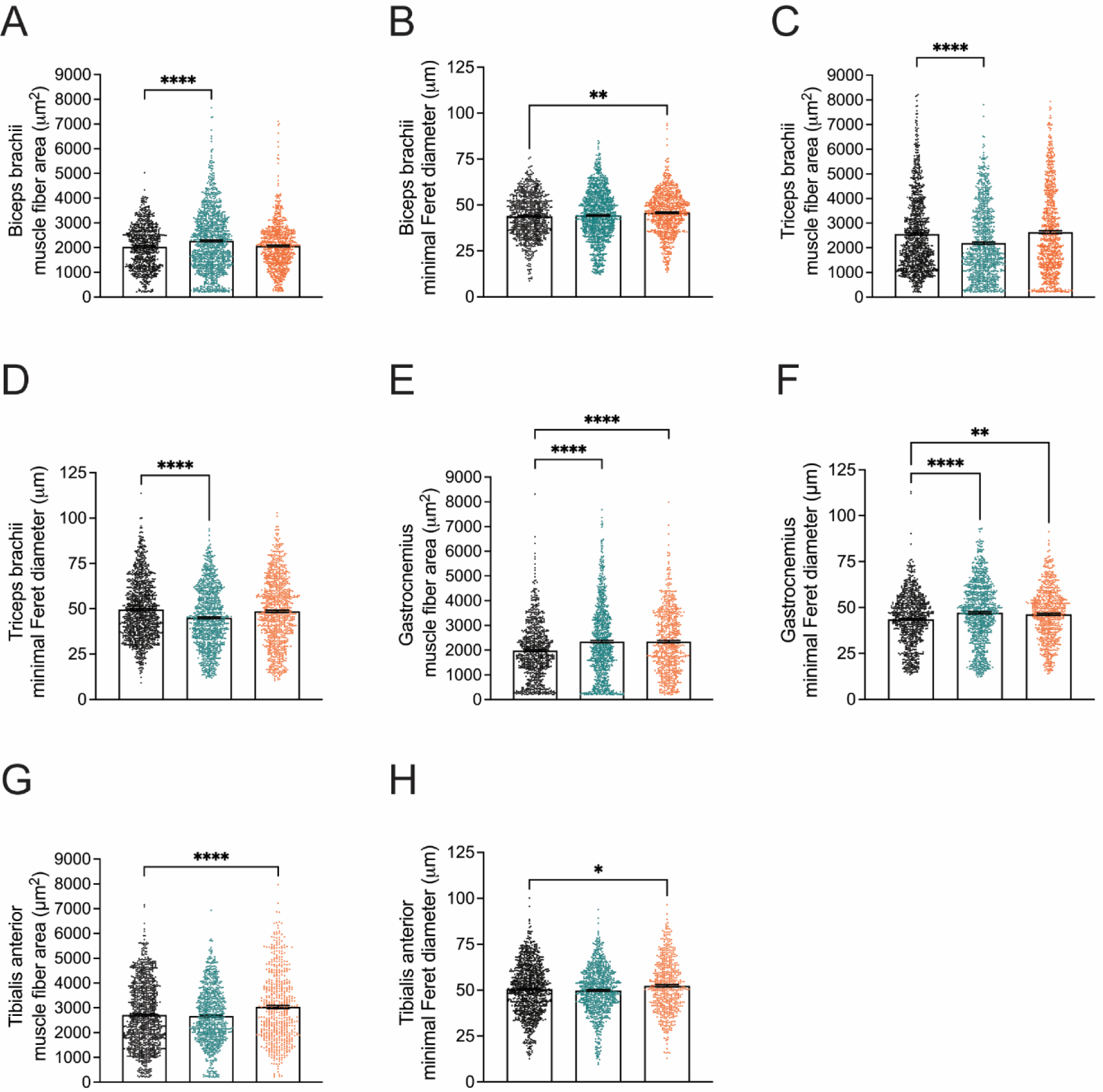
*Nefl* mutant mice have altered muscle area and diameter at twelve months. Quantification of biceps brachii, triceps brachii, gastrocnemius, and tibialis anterior muscle fiber area for *Nefl^+/+^* (black), *Nefl^+/E397K^* (teal), and *Nefl^E397K/E397K^* (orange). (A) Biceps brachii area of *Nefl^+/+^*=2032 μm^2^ (791 fibers, N=3), *Nefl^+/E397K^*=2276μm^2^ (1176 fibers, *P*<0.0001, N=5), and *Nefl^E397K/E397K^*=2069μm^2^ (781 fibers, NS, N=3). (B) Biceps brachii minimal Feret diameter of *Nefl^+/E397K^*=43.8μm (791 fibers, N=3), *Nefl^+/E397K^*=44.2μm (1176 fibers, NS, N=5), and *Nefl^E397K/E397K^*=45.8μm (781 fibers, *P*=0.0040, N=3). (C) Triceps brachii area of *Nefl^+/+^*=2557μm^2^ (1073 fibers, N=5), *Nefl^+/E397K^*=2188μm^2^ (988 fibers, *P*<0.0001, N=4), and *Nefl^E397K/E397K^*=2635μm^2^ (878 fibers, NS, N=4). (D) Triceps brachii minimal Feret diameter of *Nefl^+/+^*=49.6μm (1073 fibers, N=5), *Nefl^+/E397K^*=45.0μm (988 fibers, *P*<0.0001, N=4), and *Nefl^E397K/E397K^*=48.6μm (878 fibers, NS, N=4). (E) Gastrocnemius area of *Nefl^+/^*^E397K^ =1984μm^2^ (792 fibers, N=3), *Nefl^+/E397K^*=2349μm^2^ (918 fibers, *P*<0.0001, N=4), and *Nefl^E397K/E397K^*=2350μm^2^ (674 fibers, *P*<0.0001, N=3). (F) Gastrocnemius minimal Feret diameter of *Nefl^+/+^*=43.5μm (792 fibers, N=3), *Nefl^+/E397K^*=47.1μm (918 fibers, *P*<0.0001, N=4), and *Nefl^E397K/E397K^*=46.3μm (674 fibers, *P*=0.0013, N=3). (G) Tibialis anterior area of *Nefl^+/+^*=2716μm^2^ (1065 fibers, N=5), *Nefl^+/E397K^*=2671μm^2^ (850 fibers, NS, N=4), and *Nefl^E397K/E397K^*=3035μm^2^ (564 fibers, *P*<0.0001, N=3). (H) Tibialis anterior minimal Feret diameter of *Nefl^+/+^*=50.6μm (1065 fibers, N=5), *Nefl^+/E397K^*=49.9μm (850 fibers, NS, N=4), and *Nefl^E397K/E397K^*=52.4μm (564 fibers, *P*=0.0257, N=3). Ordinary one-way ANOVA with Dunnett multiple comparisons test was used to determine significance. N=number of mice evaluated, NS=not significant.

Changes in the muscle fiber area and diameter between *Nefl* mutants and wild type mice led us to examine the distribution of muscle fiber types (Fig. 5, Table 2). Skeletal muscle is comprised of myofibers with distinctive phenotypes that are classified based on expression of different myosin heavy chains (MyHC). Fibers expressing MyHC type I are classified as slow twitch fibers and present an oxidative metabolic type. Type I fibers express MyHC7 and are slow to contract and fatigue. MyHC type II-positive fibers are classified as fast twitch and are comprised of MyHC IIa, MyHC IIx, and MyHC IIb myofibers that express MyHC2, MyHC1, and MyHC4, respectively. Muscle fiber type composition was examined at twelve weeks and twelve months. At twelve weeks nor twelve months were significant differences in muscle fiber type composition observed in the forelimb muscles (bicep brachii and triceps brachii) or hindlimb muscles (tibialis anterior and gastrocnemius) (Fig. 5, Table 2). While significant differences were not observed, there were notable variations between wild type and *Nefl* mutant mice within muscles and fiber type composition (Table 2). At twelve weeks, there was an increase in type IIb fibers in the biceps brachii and triceps brachii of *Nefl* mutant mice when compared to wild type mice (Table 2). In contrast, the gastrocnemius showed a decrease in type IIb fibers of *Nefl* mutant mice when compared to wild type mice (Table 2). Interestingly, at twelve months there was a decrease in type IIb fibers in the biceps brachii of *Nefl* mutant mice and an increase in type IIx fibers when compared to wild type mice (Table 2). In the gastrocnemius at twelve months *Nefl*^+/E397K^ demonstrated increased IIa and IIx fibers and decreased IIb fibers when compared to wild type mice (Table 2). *Nefl*^E397K/E397K^ fiber type composition was notably different in the gastrocnemius when compared to *Nefl*^+/E397K^ mice at twelve weeks and especially at twelve months where there were decreased IIa and IIx fibers and increased IIb fibers (Table 2). The differences in muscle fiber type composition between wild type and *Nefl* mutant mice suggests that muscle function could be altered.

**Figure 5.**
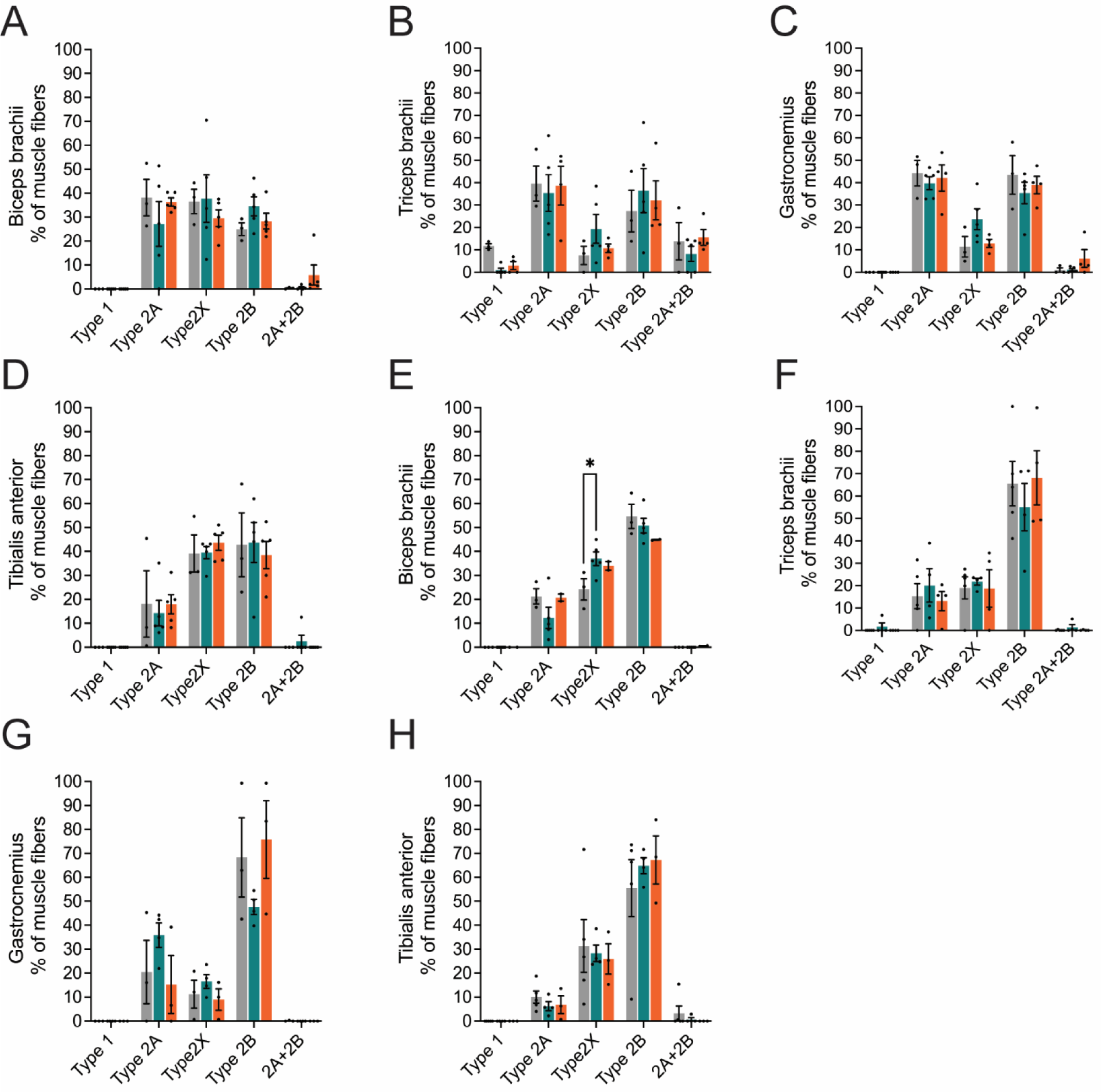
Muscle fiber type composition in the *Nelf*-E397K mouse models. Twelve weeks and twelve months cross sections of biceps brachii, triceps brachii, gastrocnemius, and tibialis anterior muscles of *Nefl^+/+^* (black), *Nefl^+/E397K^* (teal), and *Nefl^E397K/E397K^* (orange). (A) Twelve weeks biceps brachii of *Nefl^+/+^* (N=3), *Nefl^+/E397K^* (N=5), and *Nefl^E397K/E397K^* (N=5) mice. (B) Twelve weeks triceps brachii of *Nefl^+/+^* (N=3), *Nefl^+/E397K^* (N=5), and *Nefl^E397K/E397K^* (N=4) mice. (C) Twelve weeks gastrocnemius of *Nefl^+/+^* (N=3), *Nefl^+/E397K^* (N=5), and *Nefl^E397K/E397K^* (N=4) mice. (D) Twelve weeks tibialis anterior of *Nefl^+/+^* (N=3), *Nefl^+/E397K^* (N=5), and *Nefl^E397K/E397K^* (N=5) mice. (E) Twelve months biceps brachii of *Nefl^+/+^* (N=3), *Nefl^+/E397K^* (N=5), and *Nefl^E397K/E397K^* (N=2) mice. (F) Twelve months triceps brachii of *Nefl^+/+^* (N=5), *Nefl^+/E397K^* (N=4), and *Nefl^E397K/E397K^* (N=4) mice. (G) Twelve months gastrocnemius of *Nefl^+/+^* (N=3), *Nefl^+/E397K^* (N=4), and *Nefl^E397K/E397K^* (N=3) mice. (H) Twelve months tibialis anterior of *Nefl^+/+^* (N=5), *Nefl^+/E397K^* (N=4), and *Nefl^E397K/E397K^* (N=3) mice. \**P* = 0.0321. Two-way ANOVA with Šídák multiple comparisons test was used for statistical analyses. Data points/dots represent mice. N=number of mice evaluated.

**Table 2.**
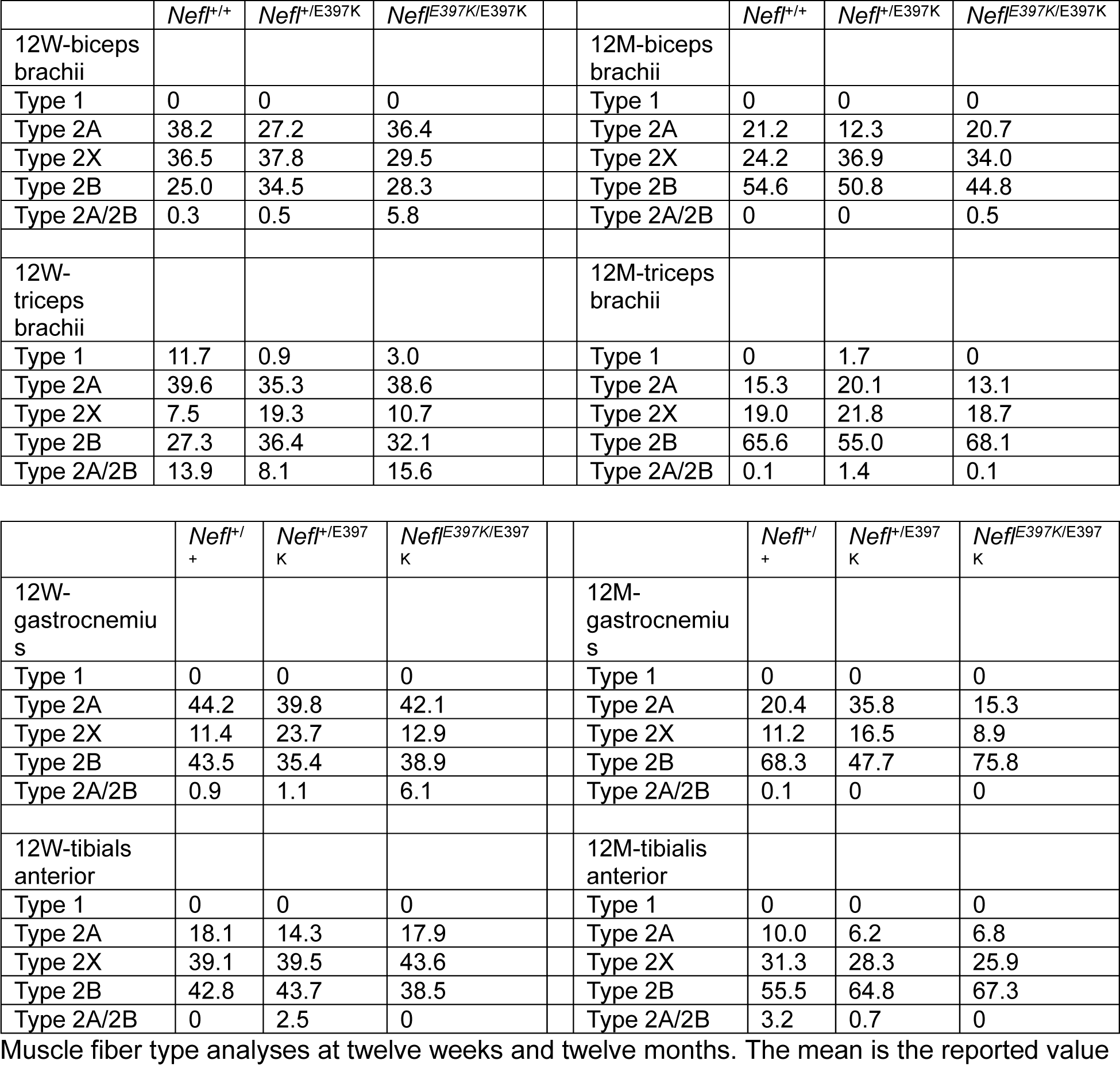

Differences in total muscle fibers, total nuclei, percent central nucleated fibers and average nuclei/fiber were initially examined in the gastrocnemius at six months of age. The gastrocnemius was chosen as it was the most affected muscle previously examined. While there were differences between *Nefl* mutants and wild type mice, these differences were not significant suggesting these parameters were not significantly altered in *Nefl* mutants (Supplementary Fig. S5A-D, I). In addition, inflammation within the biceps brachii, triceps brachii, gastrocnemius, and tibialis anterior was examined at six months of age (Supplementary Fig. S5E-I). While there were differences between *Nefl* mutants and wild type mice, these differences were not significant, suggesting muscle inflammation was not significantly increased in *Nefl* mutants.

### *Nefl* mutants show increased erratic breathing and number of apneas with increased tidal volume

A subset of CMT2E patients have reported respiratory complications, likely associated with respiratory muscle weakness. To determine if respiratory deficits were present in the *Nefl^+/E397K^* and *Nefl^E397K/E397K^* mice, whole-body plethysmography was performed to assess respiration under normal and challenge conditions at twelve weeks and twelve months (Fig. 6). Mice were evaluated under three experimental conditions: normoxia (normal, 21% O2), hypercapnia (high CO2, 7% CO2, 21% O2) and maximum challenge (high CO2, low O2, 7% CO2, 10.5% O2). Frequency (respiratory rate in breaths/minute), tidal volume (amount of air inspired), minute ventilation (rate of ventilation, frequency x tidal volume) and mean inspiratory flow (tidal volume/inspiratory time) were recorded. The total number of apneas (paused breathing) and time spent in erratic breathing (irregular breathing patterns) were also measured.

**Figure 6.**
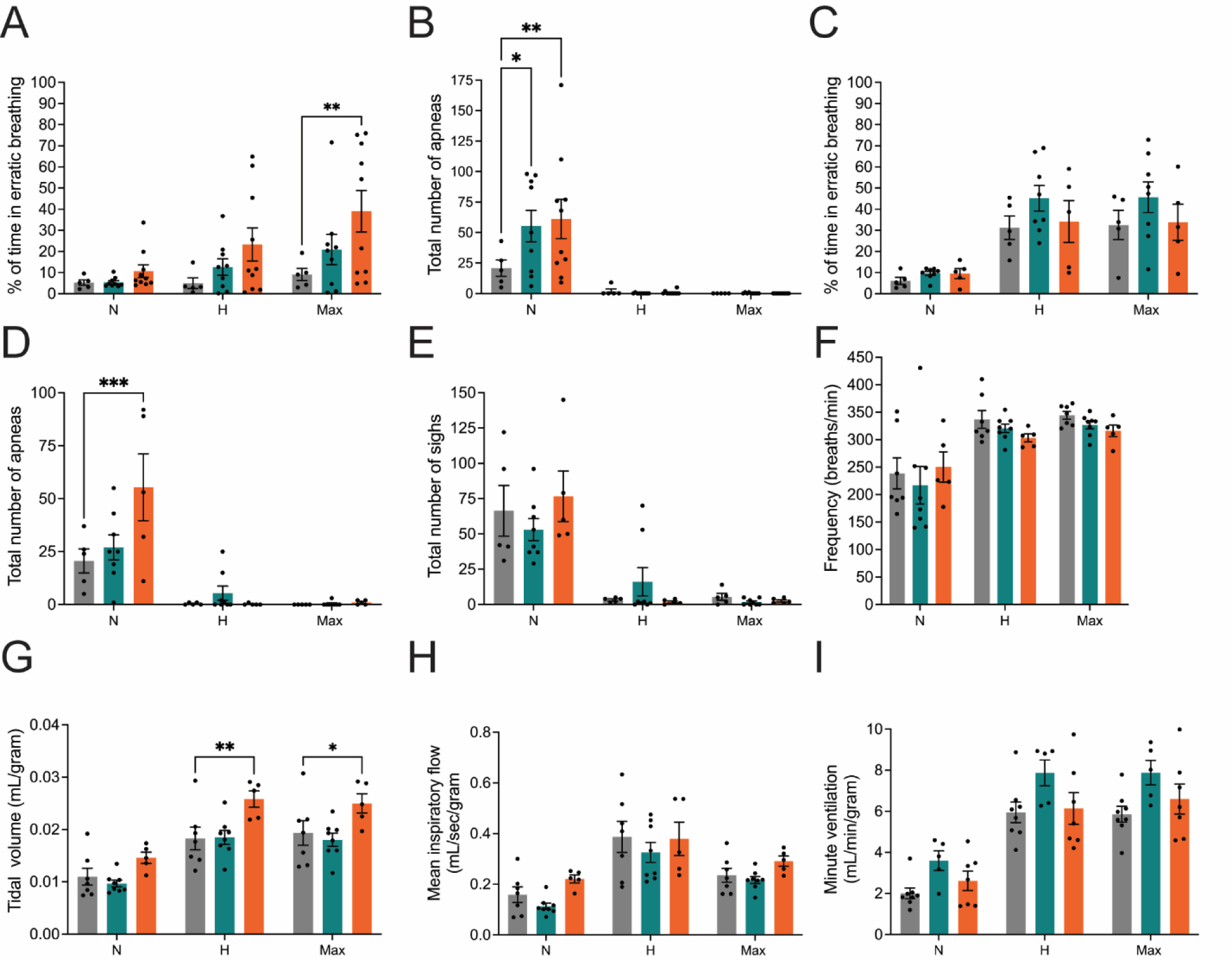
CMT2E mice presented an increased number of apneas at twelve weeks and twelve months. Quantitative whole-body plethysmography analyses of percent of time in erratic breathing, total number of apneas, total number of sighs, frequency, tidal volume, mean inspiratory flow, and minute ventilation in *Nefl^+/+^* (black), *Nefl^+/E397K^* (teal), and *Nefl^E397K/E397K^* (orange) mice. Measurements were taken under three conditions normoxia (N), hypercapnia (H), and maximum challenge (Max). At twelve weeks *Nefl^+/+^* (N=5), *Nefl^+/E397K^* (N=9) and *Nefl^E397K/E397K^* (N=10) mice were evaluated. (A) Twelve weeks percentage of time spent in erratic breathing in *Nefl^+/+^* (N=5.3, H=5.0, Max=9.1), *Nefl^+/E397K^* (N=5.4, H=12.7, Max=20.9), and *Nefl^E397K/E397K^* (N=10.7, H=23.3, Max=39.0, *P*=0.0062). (B) Twelve weeks total number of apneas in *Nefl^+/+^* (N=20.8, H=1.8, Max=0), *Nefl^+/E397K^* (N=55.2, H=0.1, Max=0.2, *P*=0.0218), and *Nefl^E397K/E397K^* (N=61.1, H=0.7, Max=0, *P*=0.0019). (C) Twelve months percentage of time spent in erratic breathing in *Nefl^+/+^* (N=6.1, H=31.2, Max=32.5), *Nefl^+/E397K^* (N=9.7, H=45.2, Max=45.7), and *Nefl^E397K/E397K^* (N=9.5, H=34.2, Max=33.8). (D) Twelve months total number of apneas in *Nefl^+/+^* (N=20.6, H=0.4, Max=0), *Nefl^+/E397K^* (N=27.0, H=5.4, Max=0.4), and *Nefl^E397K/E397K^* (N=55.4, H=0.2, Max=1.0, *P*=0.0004). (E) Twelve months total number of sighs in *Nefl^+/+^* (N=66.4, H=3.2, Max=5.4), *Nefl^+/E397K^* (N=53.0, H=16.1, Max=1.8), and *Nefl^E397K/E397K^* (N=76.6, H=1.8, Max=2.6). (F) Twelve months frequency (respiratory rate in breaths/minute) in CMT2E animals. (G) Twelve months tidal volume (amount of air inspired) *Nefl^+/+^* (N=0.011, H=0.018, Max=0.019), *Nefl^+/E397K^* (N=0.010, H=0.019, Max=0.018), and *Nefl^E397K/E397K^* (N=0.015, H=0.026, Max=0.025, \*\**P*=0.0062, \**P*=0.0452). (H) Twelve months mean inspiratory flow (tidal volume/inspiratory time). (I) Twelve months minute ventilation (rate of ventilation, frequency x tidal volume). Two-way ANOVA with Dunnett multiple comparison test was used to determine significance. Data points/dots represent mice. N=number of mice evaluated.

At twelve weeks there were no significant differences between *Nefl* mutant mice and wild type mice in frequency, tidal volume, minute ventilation and mean inspiratory flow during normoxia, hypercapnia or max challenge (not shown). However, there were differences between *Nefl* mutant and wild type mice at twelve-weeks in the percent time spent in erratic breathing and the total number of apneas (Fig. 6A, B). Under hypercapnia conditions, wild type mice spent 5.0% of the recorded time in erratic breathing while *Nefl*^+/E397K^ mice spent 12.7% and *Nefl*^E397K/E397K^ mice spent 23.3% time in erratic breathing. Erratic breathing was increased further under maximum respiratory challenge conditions where wild type mice spent 9.1% of the recorded time in erratic breathing while *Nefl*^+/E397K^ mice spent 21.0% and *Nefl*^E397K/E397K^ mice spent 39.0% time in erratic breathing (Fig. 6A). In *Nefl* mutant mice there was more variation between mice suggesting some mice had deficits more than others. When apneas were evaluated at twelve weeks differences were noted only under normoxia conditions (Fig. 6B). Wild type mice recorded 20.8 apneas under normoxia conditions while *Nefl*^+/E397K^ mice recorded 55.2 apneas and *Nefl*^E397K/E397K^ mice recorded 61.1 apneas. Concerning apneas, there was more variation between *Nefl* mutant mice suggesting some mice had deficits more than others.

At twelve months, *Nefl* mutant mice continued to show increased number of apneas under normoxia conditions (Fig. 6D). Wild type mice recorded 20.6 apneas under normoxia conditions while *Nefl*^+/E397K^ mice recorded 27.0 apneas and *Nefl*^E397K/E397K^ mice recorded 55.4 apneas. There was more variation between *Nefl* mutant mice suggesting some mice had deficits more than others with *Nefl*^E397K/E397K^ mice experiencing the most apneas. Interestingly, *Nefl*^E397K/E397K^ mice demonstrated significantly increased tidal volume under hypercapnia and maximum challenge conditions (Fig. 6G). Tidal volume under hypercapnia conditions was 0.018 mL/gram for wild type mice while *Nefl*^+/E397K^ mice recorded a tidal volume of 0.019 mL/gram and *Nefl*^E397K/E397K^ mice 0.026 mL/gram. Tidal volume under maximum challenge conditions was 0.019 mL/gram for wild type mice while *Nefl*^+/E397K^ mice recorded a tidal volume of 0.018 mL/gram and *Nefl*^E397K/E397K^ mice 0.025 mL/gram. The increased tidal volume suggests some compensatory mechanism in *Nefl*^E397K/E397K^ mice under challenged respiratory conditions.

## Discussion

Rodent models of disease are essential for translational research as they provide valuable insights into disease mechanisms and therapeutic strategies. While no model can fully capture the complexity of human biology, their value lies in their ability to mimic critical elements of a disease rather than recreate it entirely, allowing researchers to test hypotheses and refine therapies before clinical trials. Several mouse models have been developed to study CMT2E, each with its own strengths and limitations. In this report, we demonstrate that the novel *Nefl^+/E397K^* and *Nefl^E397K/E397K^* mouse models of CMT2E present with important, progressive features of this disease, including motor function deficits, and skeletal muscle fiber pathology. Additionally, these mice also show respiratory apneas and erratic breathing differences similar to those reported in CMT patients. This model is an important addition to the CMT2E collection of animal models as it has a robust and early (at least by P21) phenotype that is progressive, but not so severe that the animals cannot be studied over an extended period (at least up to P360).

When evaluating motor function assessments, *Nefl*^E397K/E397K^ mice often performed worse than *Nefl*^+/lE397K^ mice. Consistent with the CMT2E patient population, variability among *Nefl* mutant was noted and this variability often masked the true significance of the motor function deficits. Interestingly, deficiencies were observed earliest in assessments that required the most motor coordination (rotarod and dowel rod) and these deficits became progressively worse. The deterioration of motor control was also observed in how *Nefl* mutant mice completed the motor function assessments. Gait was reduced while the number of tail grabs and foot slips increased. Tail and foot positioning while traversing the dowel rod also indicated that *Nefl* mutant mice experienced motor control deficits. Catwalk analyses provided further insight into the gait disturbances observed in the hind paws maximal contact area and print width as well as the left and right print positions. These motor coordination deficits were consistent with the electrophysiological findings observed at three weeks through twelve months (companion manuscript).

*Nefl* mutant mice demonstrated dynamic changes in muscle fiber area and diameter as disease progressed that varied not only between forelimbs and hindlimbs but within forelimbs and hindlimbs. The forelimb biceps brachii and hindlimb gastrocnemius demonstrated the most significantly changes in muscle fibers that could be attributed to the muscle composition with more fast twitch muscle fibers found within these muscles when compared to the triceps brachii and tibialis anterior. At twelve weeks, the biceps and gastrocnemius muscles showed significant atrophy in *Nefl*^+/lE397K^ and *Nefl*^E397K/E397K^ mice. In contrast, at twelve months hypertrophy was observed in the biceps and gastrocnemius of *Nefl*^+/lE397K^ and *Nefl*^E397K/E397K^ mice. Atrophy of the biceps and gastrocnemius is likely attributed to the significant axonal degeneration observed and is supported by the electrophysiology findings (companion manuscript). Additionally, neuromuscular junction denervation was increased in *Nefl* mutants when compared to three weeks (companion manuscript). While some inflammation was observed within the gastrocnemius, inflammation was likely not the primary contributor to hypertrophy observed but rather some compensatory mechanisms. Axonal area and diameter and the number of axons/field all increased from six months to twelve months supporting compensatory mechanisms at play (companion manuscript). In contrast to the biceps brachii and gastrocnemius, the triceps brachii showed hypertrophy at twelve weeks followed by atrophy at twelve months in *Nefl*^+/E397K^ mice with no significant changes observed in *Nefl*^E397K/E397K^ mice. These differences are likely attributed to differences in muscle composition and function or differences in axonal degeneration. It is unclear why *Nefl*^+/E397K^ mice showed alterations in the triceps while *Nefl*^E397K/E397K^ mice did not.

Muscle fiber type within the biceps brachii, triceps brachii, gastrocnemius and tibialis anterior demonstrated alterations between wild type and *Nefl* mutant mice and throughout the disease. Changes in fiber type composition could lead to changes in muscle function, muscle weakness and reduced motor function observed in the *Nefl* mutant mice. At twelve weeks, the biceps brachii (slightly higher type II fibers) showed increased type IIb fibers in both *Nefl* mutants while having decreased muscle fiber area and diameter within the biceps. This is interesting in that type IIb fibers are larger fibers when compared to type IIa and IIx fibers suggesting there are more, smaller type IIb fibers. At twelve weeks, the distribution of biceps brachii muscle fibers shifted to more, smaller fibers in both *Nefl* mutants. At twelve months, *Nefl*^+/E397K^ and *Nefl*^E397K/E397K^ mice showed decreased type IIb fibers and a switch to increased type IIx fibers in the biceps brachii. This alteration could be attributed to a change in motor units as a compensatory mechanism towards regeneration as muscle fiber area and diameter were either similar to or greater in the *Nefl* mutants when compared to wild type. The gastrocnemius, a primarily fast twitch muscle with a high proportion of type IIb fibers, showed decreased muscle fiber area and diameter as well as decreased type IIa and IIb fibers at twelve weeks in both *Nefl* mutants consistent with the decreased distribution of larger muscle fibers and atrophy. At twelve months, increased muscle fiber area and diameter were apparent and there were shifts in the proportions of type II muscle fibers. *Nefl*^+/E397K^ mice had decreased type IIb fibers and increased type IIa and IIx fibers while *Nefl*^E397K/E397K^ mice had increased type IIb fibers and decreased type IIa and IIx fibers. These results along with the increased sciatic axon area, diameter and axon numbers/field suggest a regenerative process taking place at twelve months within the gastrocnemius.

We report the first respiratory deficiencies in CMT2E model mice, consistent with the observation that a portion of CMT patients present pulmonary and sleep apneas (22). Consistent with the patient population, there was more variability in respiratory measurements between *Nefl* mutant mice within a cohort. Two different types of apneas have been identified in CMT patients, central sleep apneas and obstructive sleep apneas. Central sleep apneas may by associated with chemosensory and diaphragm dysfunction. Obstructive sleep apneas are caused by physical obstruction during breathing, and this obstruction can be an anatomical abnormality (23). At twelve weeks, *Nefl^+/E397K^* and *Nefl^E397K/E397K^* showed an increase in apneas under normoxia conditions that persisted through twelve months. At twelve months, increased tidal volume was also observed under hypercapnia and maximal challenge conditions indicative of reduced forced vital capacity, similar to CMT patients (24). These models provide a novel experimental context to address important disease features as well as provide critical clinical features for future drug development efforts.

It will be important to leverage these *Nefl* models appropriately based upon the experimental questions. For example, for a longitudinal study to examine potential biomarkers, the *Nefl*^+/E397K^ may be more appropriate. For a gene therapy study, the *Nefl^E397K/E397K^* model may be more suitable even though the “true” disease state is typically hemizygous simply because of the greater degree of disease severity and because of the age of onset. Collectively, these models further enhance our understanding of disease progression and provide important experimental contexts for future therapeutic development.

## Materials and Methods

### Animals

All experimental procedures were approved by the University of Missouri Animal Care and Use committee and were performed according to the guidelines set forth in the Guide for the Use and Care of Laboratory Animals. *Nefl* mice on a C57BL/6 background were generated using CRISPR technology at the University of Missouri Animal Modeling Core and *Nefl^+/E397K^* were used as breeders. *Nefl^+/E397K^, Nefl^E397K/E397K^,* and wild type male and female littermates were assessed from postnatal day (P) 1 to P360.

### Genotyping

Genotyping of neonatal pups was performed at ∼P1. Genomic DNA isolation was performed using a protocol from Jacksons labs. THE C57BL/6J-*Nefl^E397K^* line was genotyped with a high-resolution melt analysis was performed using an EvaGreen qPCR precision melt supermix (Bio-Rad) and the primers 5’-CTTCATTCCCTCTCCACCAG-3’ and 5’-CACGCTGGTGAAACTGAG-3’. The high-resolution melt PCR conditions were 95°C denaturing for 2:00 minutes followed by 39 cycles of 95°C denaturing for 10 seconds, 60°C annealing for 30 seconds and 72°C extension for 30 seconds, 95°C for 30 seconds, 60°C for 1:00 minute, and melt curve 65°C to 95°C in increments of 0.2 every 10 seconds. Melting temperature for the wild type, *Nefl^+/E397K^*, and *Nefl^E397K/E397K^* were 78.5 ± 0.03°C, 77.7 ± 0.03°C, and 77.2 ± 0.06°C respectively.

### Motor function assessments

Hindlimb splay (HLS) was implemented to assess hindlimb function and was initiated at P7 as previously reported. In short, mice were held ∼0.5” from the base of the tail over the wire top of the cage. The relative distance between the splayed limbs was recorded as a qualitative measurement, a score of three was assigned to a fully splayed hindlimb, score of two was assigned when the hindlimbs were parallel to the body, score of one was assigned when the hindlimbs were towards midline, and score of zero when the hindlimbs were in contact or clasping.

The time to right (TTR) assay assessed core muscle strength daily from P8 until P30. Each pup was placed on its backside and the time (seconds) it took to turn over and stabilize on all four paws was recorded.

Grip strength assays assessed skeletal muscle strength in the forelimb and hindlimbs. Mice were gently pulled by the back by their tail ensuring each mouse gripped the grid and the torso remained horizontal. The maximal grip strength value was recorded (BIO-GS4, Bioseb). All limb grip strength values were averaged across three trials per assessment. For hindlimb grip strength, mice were scruffed and their hind paws were placed on one rung of the grate, which triggers an involuntary grip reflex. Mice were gently pulled with consistent force across bottom three rungs and the value recorded and averaged across three trials. Mice were trained for three days before data collection at P35. Grip strength assessments were conducted weekly starting at P35 to P84, then at six and twelve months.

Rotarod (ITC Rotarod Series 8, IITC Life Science) assessments were performed as previously reported. The time the mouse walked forward on the rotating rod was measured in seconds for three trials/assessment and then averaged. The initial speed was 4 rpm with a gradual increase to 40 rpm over 150 seconds. Mice were trained for three days before data collection at P28. Assessments were conducted weekly starting at P35 to P77 then monthly from P120 up to 6 months. Longitudinal rotarod assessments from P35-P180 were collected.

Gait abnormalities were measured using the CatWalk (Noldus) automated system to measure individual footprint-based parameters. On each paw measurements of stride length (distance between successive placements of the same paw), print positions (position of the hind paws relative to the previous position of the fore paw), box width (width of the complete footprint), and box length (indicates the length of the complete footprint) were taken. Mice were trained at P28, and two trials were recorded weekly from P35 up to P84 and then monthly from P90 to P360.

Dowel rod (22 mm diameter) testing was used to assess coordination, balance, and motor function. Two trials were recorded for each mouse and analyses were performed in a blinded manner from the recorded videos. Mice were placed on a suspended dowel (26 inches in length) and given twenty seconds to cross from one end of the dowel to the other. Mice were assessed for tail grabs, foot slips, falls, and the time to transverse the rod. Balance was assessed by scoring tail positioning from 1 to 3, where 1 indicates a tail used to balance and support, a score of 2 indicates a tail parallel to the dowel, and score of 3 indicates a tail up without any visible balance problems. Foot slips and tail grabs were quantified on each video and reported as the average. Mice were trained at P30, and average of two trials were recorded weekly from P35 up to P84 and then at P180 and P360.

### Skeletal muscle immunohistochemistry

Fresh tissue was embedded in OCT and flash frozen in liquid nitrogen-chilled 2-methyl-2-butene. 10 μm sections were co-stained with anti-Laminin primary antibody (1:200; catalog: L9393; Millipore Sigma), myosin heavy chain (MyHC) type I (1:10, catalog: SC-71-s, Iowa Developmental Studies Hybridoma Bank), MyHC type IIa (1:50, catalog: BA-D5-s, Iowa Developmental Studies Hybridoma Bank), MyHC type IIb (1:5, catalog: BF-F3-s, Iowa Developmental Studies Hybridoma Bank) as previously reported (25). Secondary antibody for laminin was Goat anti-Rabbit, Alexa Fluor^TM^ 647 (1:500, catalog: A21244, ThermoFisher Scientific), (MyHC) type I Goat anti-Mouse, Alexa Fluor^TM^ 350 (1:500, catalog: A21140, ThermoFisher Scientific), MyHC type IIa Goat anti-Mouse, Alexa Fluor^TM^ 555 (1:500, catalog: A21127, ThermoFisher Scientific), and MyHC type IIb Goat anti-Mouse, Alexa Fluor^TM^ 488 (1:500, catalog: A21042, ThermoFisher Scientific). A random selected field at 10X magnification was obtained using a Leica DM5500 B fluorescent microscope (Leica Microsystem Inc.). Muscle fiber area, minimal Feret diameter, and muscle fiber type composition were analyzed in a blinded manner using SMASH, semi-automatic muscle analysis using segmentation of histology (26).

### Hematoxylin and eosin staining

Fresh tissues were harvested, embedded in in OCT media, and cryosectioned at 10 μm. Hematoxylin and eosin staining was performed as previously described (27). Tissue slides were fixed with acetic acid alcohol for 5 minutes, rinsed in tap water twice for 15 seconds and then hematoxylin (catalog: 411165000, ThermoFisher Scientific) stained for 1.5 minutes. Slides were then rinsed in ammonia water for 15 seconds, dehydrated in 95% ethanol twice for 15 seconds and eosin Y (catalog: 152885000, ThermoFisher Scientific) stained for 15 seconds. Finally, slides were rinsed twice for 15 seconds in 95% ethanol, 100% ethanol, and xylene. Tissues were covered with Permount mounting medium (catalog: SP15-500, ThermoFisher Scientific). Three individual images at were acquired at 20x magnification using a Leica DM5500 B microscope (Leica Microsystem Inc.) and analyzed blindly. Quantification of all peripheral and central nuclei were performed in ImageJ (NIH) and average percentages of the three images were calculated. The freehand selection tool on ImageJ was used to marked inflamed areas and percentages were calculated based of the total area of the image, and average percentages of the three images were presented.

### Plethysmography

Mice were placed in a whole-body plethysmography chamber (Data Sciences International/Harvard Bioscience, Holliston, MA) to allow for quantitative measurement of ventilation through different gas concentrations (gas concentrations controlled by a gas mixer, CWe, Inc., Ardmore, PA). Ventilation was assessed at twelve weeks and twelve months in wild type and CMT2E mice. The mice were acclimated to the chamber while breathing room air (21% O2, balance N2) for thirty minutes before ventilatory measurements were recorded for baseline conditions (still under 21% O^2^, balance N^2^) for an additional thirty minutes. Ventilatory measurements were then made during exposure to a hypoxic + hypercapnic gas mixture (10.5% O2, 7% CO2) for five minutes. The pressure calibration signal, ambient and chamber pressures, and mouse body mass were used to calculate the respiratory frequency (f), tidal volume (V^T^), minute ventilation (VE), mean inspiratory flow (V^T^/T^i^), using Buxco FinePointe Software (Data Sciences International/Harvard Bioscience, Holliston, MA). In addition, the apnea detection function within FinePointe software was used to identify the percentage of erratic breathing (defined as any breathing that was not classified as a normal breath, sigh, apnea or sniff; Data Sciences International/Harvard Bioscience, Holliston, MA), and apneas that were defined as the absence of at least two inspirations (i.e. a pause in breathing 2x the normalized breath duration threshold). V^T^, VE and V^T^/T^i^ are reported normalized to mouse body weight (per gram). Apnea calculations were based on the minimum apnea duration based on the frequency and calculated as (60/frequency) x 2. Parameters for apneas data analysis were the following: scoring interval: 00:00:30, breathing baseline interval: 00:00:10, combined thresholds using the minimum apnea duration and percent increase in duration of 200 (twelve weeks) or 300 (twelve months), normalized V^T^ threshold with a 150% increase in V^T^ minimum sniff rate of 300, and maximum sniff volume (mL) of 0.06. Data were rejected if there was evidence of pressure fluctuations caused by gross body movements (28).

## Funding

This work was supported by the Charcot-Marie-Tooth Research Foundation (grant 00070082). DPL was supported by the MU Life Sciences Fellowship, National Institutes of Health Training grant T32 GM008396 and a Southeastern Conference (SEC) Scholar fellowship. ZA and MA were funded by the IMSD/MARC program National Institutes of Health Training grant T34 GM136493. ZA was also funded by the Discovery Fellows program at MU. AAS was funded by the MU Cherng Summer Scholars program.

## Acknowledgements

We acknowledge the MU Animal Modeling Core, MU Genomics Technology Core, and MU Advanced Light Microscopy Core for assistance with these studies. The antibodies used for muscle fiber type characterization were obtained from the Developmental Studies Hybridoma Bank, created by the NICHD of the NIH and maintained at The University of Iowa, Department of Biology, Iowa City, IA 52242.

## Conflict of Interest Statement

<u>Ownership</u>: CLL is co-founder and chief scientific officer of Shift Pharmaceuticals. MAL is associated with Shift by family relation. <u>Income</u>: CLL has received more than $10,000 in income per annum from Shift Pharmaceuticals. <u>Research support</u>: Research in the CLL and MAL labs have been supported by sub-awards from Shift Pharmaceuticals (as part of grants from the DOD, CMT Research Foundation, and the NIH). <u>Intellectual property</u>: an invention disclosure has been submitted covering the intellectual property related to this work.

## Author Contributions

DPL: designed research studies, conducted experiments, acquired data, analyzed data, prepared manuscript, edited manuscript

AS: conducted experiments, acquired data, analyzed data

FLT: conducted experiments, acquired data, analyzed data

RM: conducted experiments, acquired data, analyzed data

ZA: conducted experiments, acquired data, analyzed data

MA: conducted experiments, acquired data, analyzed data

CS: conducted experiments, acquired data

NN: analyzed data

MAL: conducted experiments, acquired data, analyzed data, edited manuscript

CLL: designed research studies, analyzed data, edited manuscript

---

**Supplementary Figure S1A-H.**
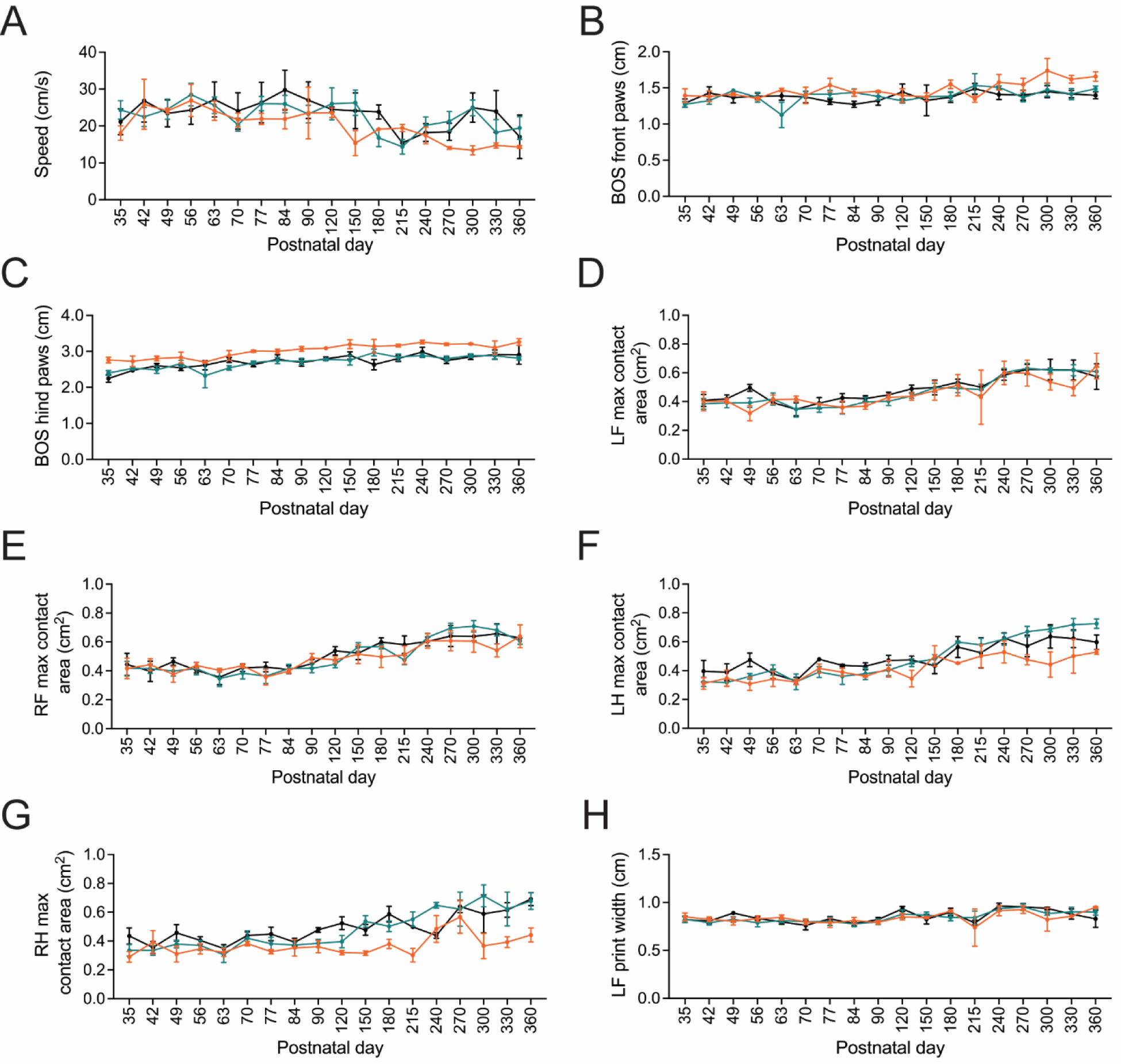
CatWalk assessments in *Nefl* mutant mice show differences. Longitudinal CatWalk assessment was performed in *Nefl^+/+^* (black), *Nefl^+/E397K^* (teal), and *Nefl^E397K/E397K^* (orange) mice to evaluate individual footprint-based parameters. (A) Speed measured in centimeters per second (cm/s). (B) Base of support (BOS) of front paws in centimeters (cm). (C) BOS of hind paws in centimeters (cm). (D) Left front (LF) maximal contact area. Maximal contact indicates the moment during a stance phase for which the footprint is largest recorded in centimeters squared. (E) Right front (RF) maximal contact area recorded in centimeters squared. (F) Left hind (LH) maximal contact area recorded in centimeters squared. (G) Right hind (RH) maximal contact area recorded in centimeters squared. (H) Left front (LF) print width in centimeters. Print width Indicates the width of the complete footprint. Mixed-effect analysis with a Bonferroni’s multiple comparison test and two-way ANOVA and repeated measures one-way ANOVA with Dunnett multiple comparison test were used to determine significance.

**Supplementary Figure S1I-M.**
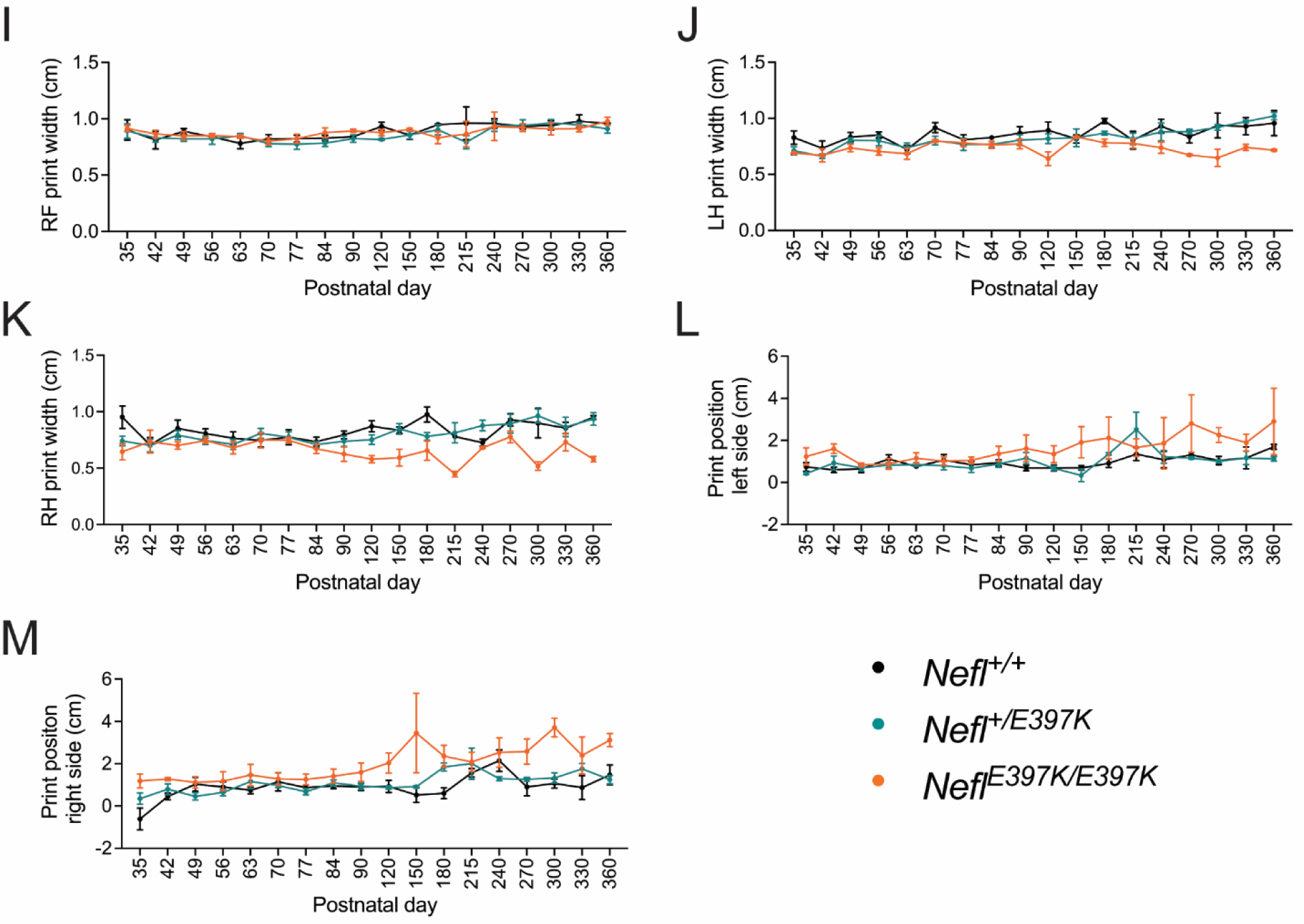
CatWalk assessments in *Nefl* mutant mice show differences. Longitudinal CatWalk assessment was performed in *Nefl^+/+^* (black), *Nefl^+/E397K^* (teal), and *Nefl^E397K/E397K^* (orange) mice to evaluate individual footprint-based parameters (I) Right front (RF) print width in centimeters. (J) Left hind (LH) print width in centimeters. (K) Right hind (RH) print width in centimeters. (L) Print position of left side paws. Print position indicates the position of the hind paw relative to the previous position of the fore paw. (M) Print position of right side paws. Mixed-effect analysis with a Bonferroni’s multiple comparison test and two-way ANOVA and repeated measures one-way ANOVA with Dunnett multiple comparison test were used to determine significance.

**Supplementary Figure S2.**
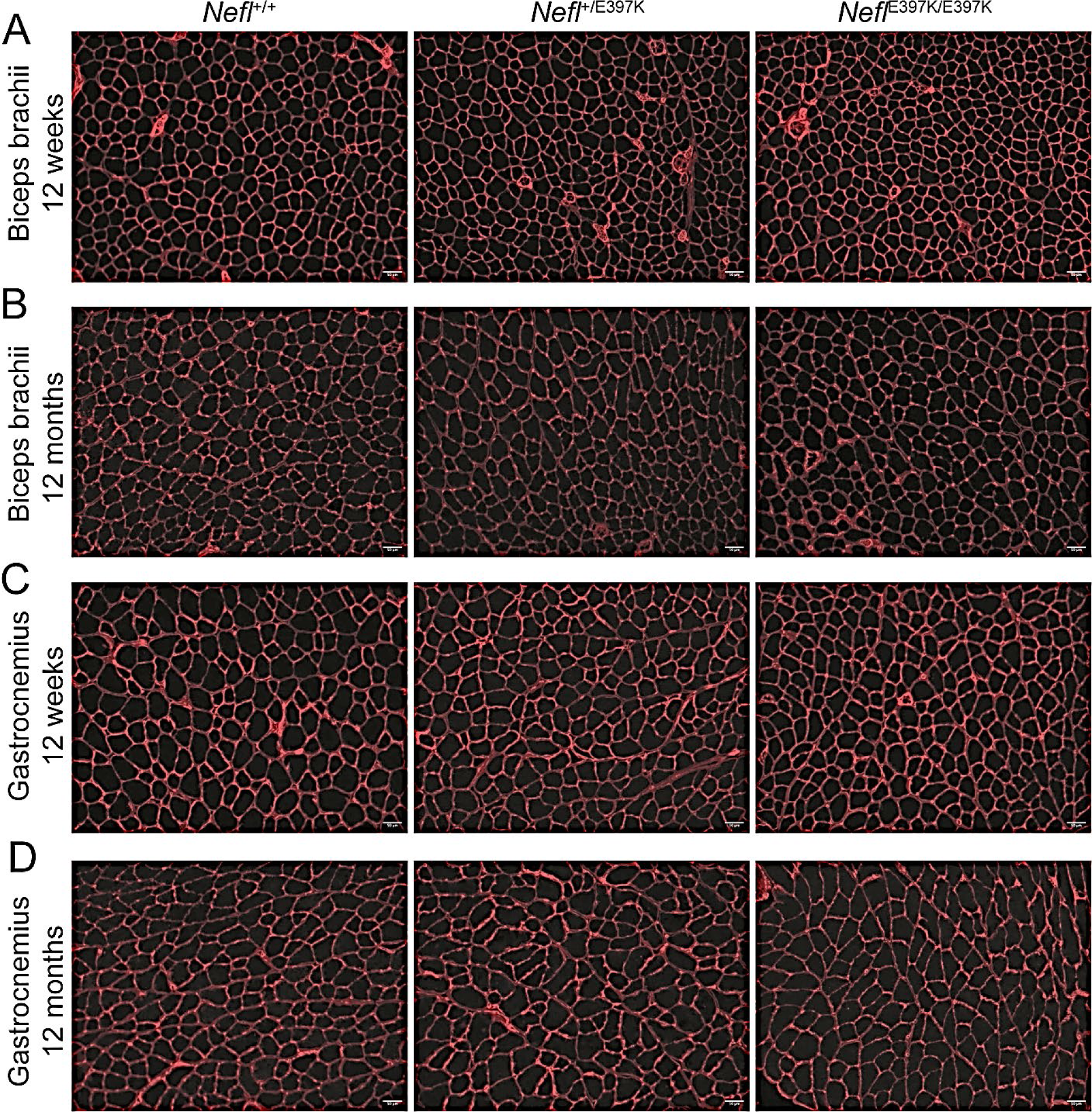
Representative muscle fiber images. Representative cross-sectional images of muscle fibers of biceps brachii and gastrocnemius muscles at twelve weeks and twelve months of *Nefl^+/+^*, *Nefl^+/E397K^*, and *Nefl^E397K/E397K^* mice immunolabeled with laminin. (A) Biceps brachii cross sections of twelve weeks old mice. (B) Biceps brachii cross sections of twelve months mice. (C) Gastrocnemius cross sections of twelve weeks old mice. (D) Gastrocnemius cross sections of twelve months old mice.

**Supplementary Figure S3.**
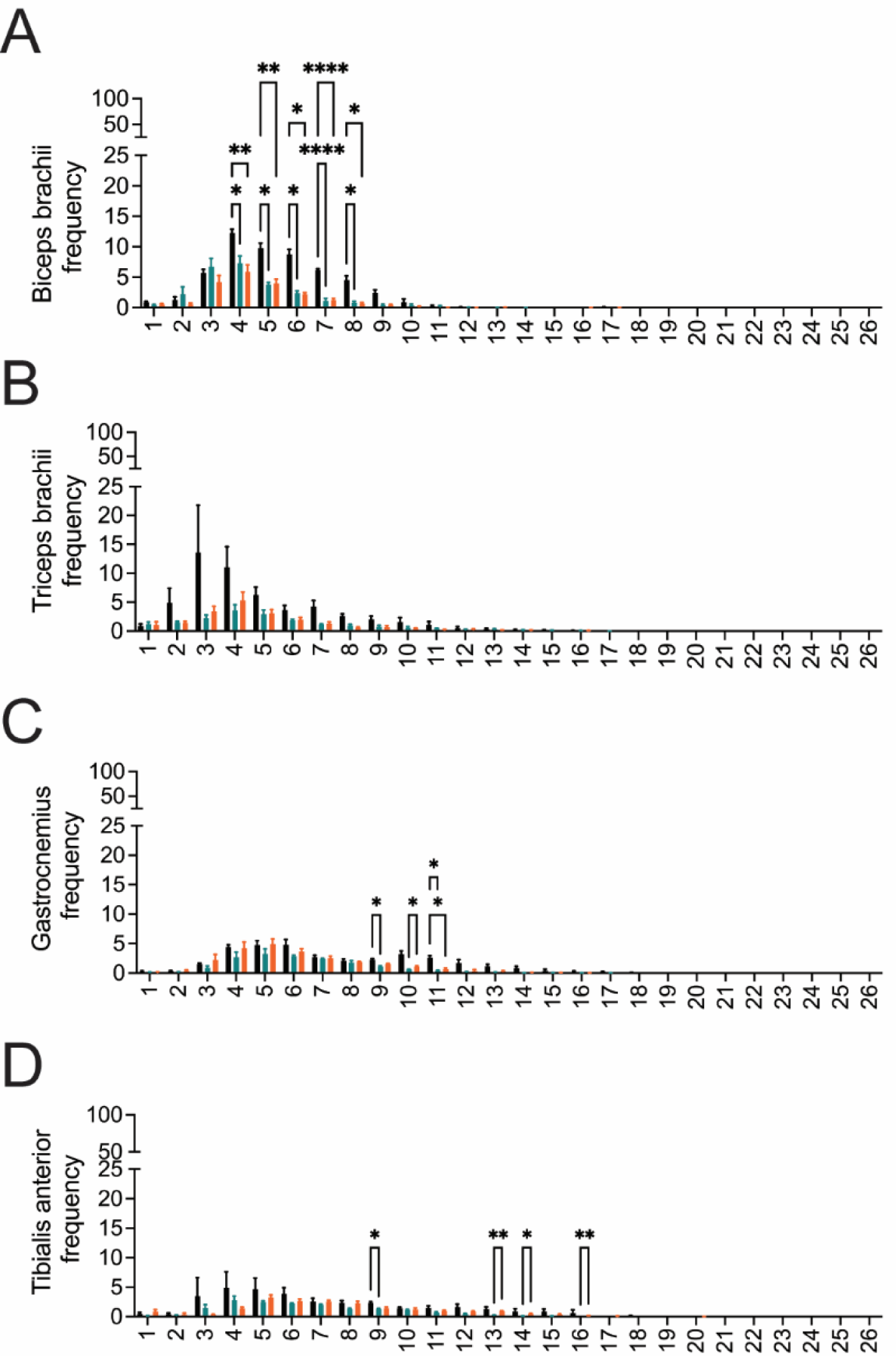
*Nefl* mutants show changes in the distribution of muscle fiber size. Muscle fiber area distribution in the forelimbs and hindlimb muscles of *Nefl^+/+^* (black), *Nefl^+/E397K^* (teal), and *Nefl^E397K/E397K^* (orange) mice at twelve weeks. Cross-sections of individual muscle fibers of biceps brachii, triceps brachii, gastrocnemius, and tibialis anterior muscles were binned by area into 400 μm^2^ intervals and expressed as percentage of muscle fibers. (A) Biceps brachii muscle fiber area distribution (\**P*=0.0434-0.0101, \*\**P*=0.0076, 0.0072, \*\*\*\**P*<0.0001). (B) Triceps brachii muscle fiber area distribution. (C) Gastrocnemius muscle fiber area distribution (*\*P*=0.0362-0.222). (D) Tibialis anterior muscle fiber area distribution (\**P*=0.0438, 0.0435, \*\**P*=0.0022, 0.0028). Distribution in μm^2^ 1=0-400, 2=400-800, 3=800-1200, 4=1200-1600, 5=1600-2000, 6=2000-2400, 7=2400-2800, 8=2800-3200, 9=3200-3600, 10=3600-4000, 11=4000-4400, 12=4400-4800, 13=4800-5200, 14=5200-5600, 15=5600-6000, 16=6000-6400, 17=6400-6800, 18=6800-7200, 19=7200-7600, 20=7600-8000, 21=8000-8400, 22=8400-8800, 23=8800-9200, 24=9200-9600, 25=9600-10000, 26=>10000. Mixed-effect analysis with Tukey’s multiple comparison test were used to determine significance.

**Supplementary Figure S4.**
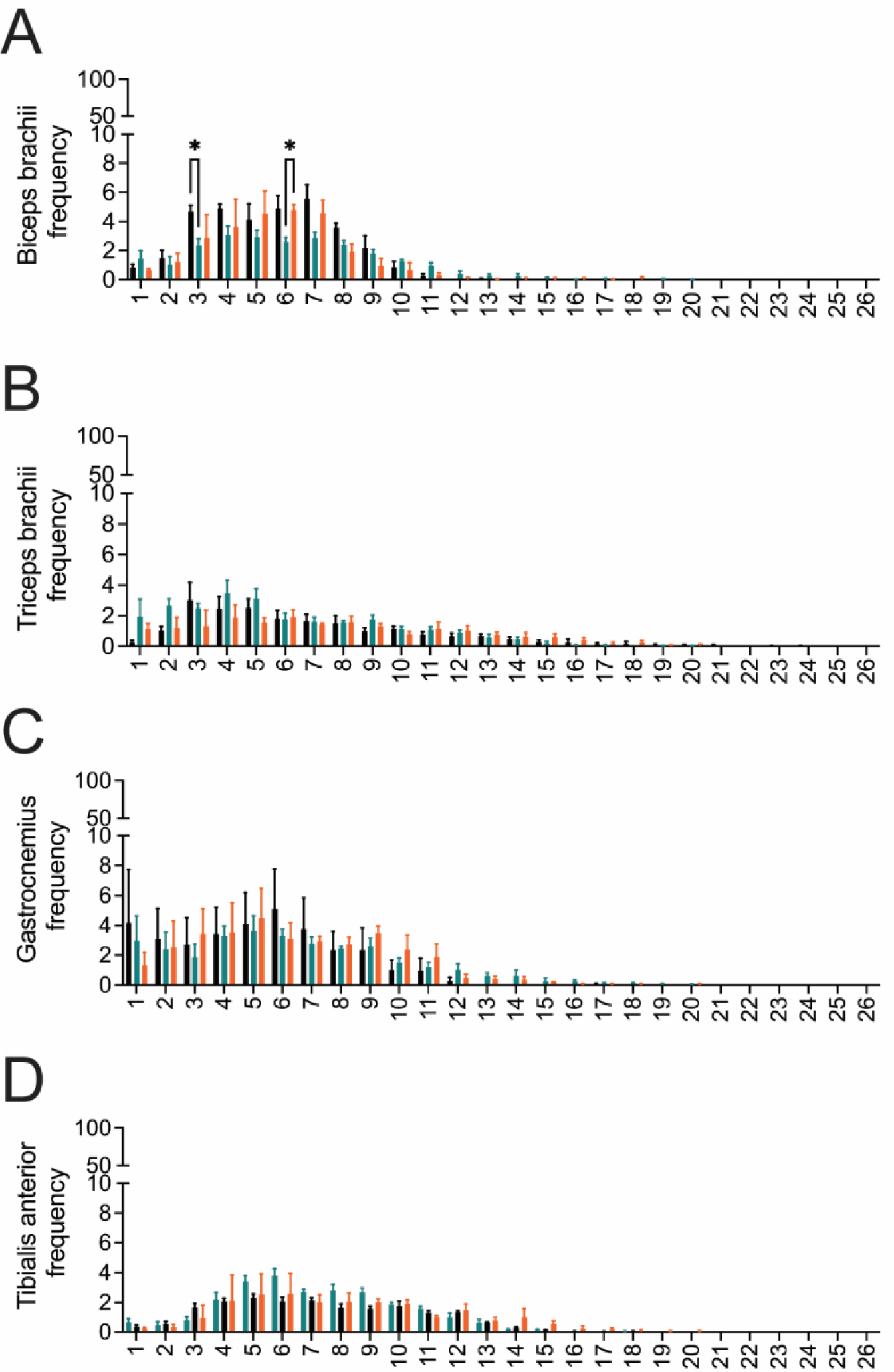
The distribution of muscle fiber size at twelve months. Muscle fiber area distribution in the forelimbs and hindlimb muscles of *Nefl^+/+^* (black), *Nefl^+/E397K^* (teal), and *Nefl^E397K/E397K^* (orange) mice at twelve months. Cross-sections of individual muscle fibers of biceps brachii, triceps brachii, gastrocnemius, and tibialis anterior muscles were binned by area into 400 μm^2^ intervals and expressed as percentage of muscle fibers. (A) Biceps brachii muscle fiber area distribution (\**P*=0.0283, 0.0145). (B) Triceps brachii muscle fiber area distribution. (C) Gastrocnemius muscle fiber area distribution. (D) Tibialis anterior muscle fiber area distribution. Distribution in μm^2^ 1=0-400, 2=400-800, 3=800-1200, 4=1200-1600, 5=1600-2000, 6=2000-2400, 7=2400-2800, 8=2800-3200, 9=3200-3600, 10=3600-4000, 11=4000-4400, 12=4400-4800, 13=4800-5200, 14=5200-5600, 15=5600-6000, 16=6000-6400, 17=6400-6800, 18=6800-7200, 19=7200-7600, 20=7600-8000, 21=8000-8400, 22=8400-8800, 23=8800-9200, 24=9200-9600, 25=9600-10000, 26=>10000. Mixed-effect analysis with Tukey’s multiple comparison test were used to determine significance.

**Supplementary Figure S5.**
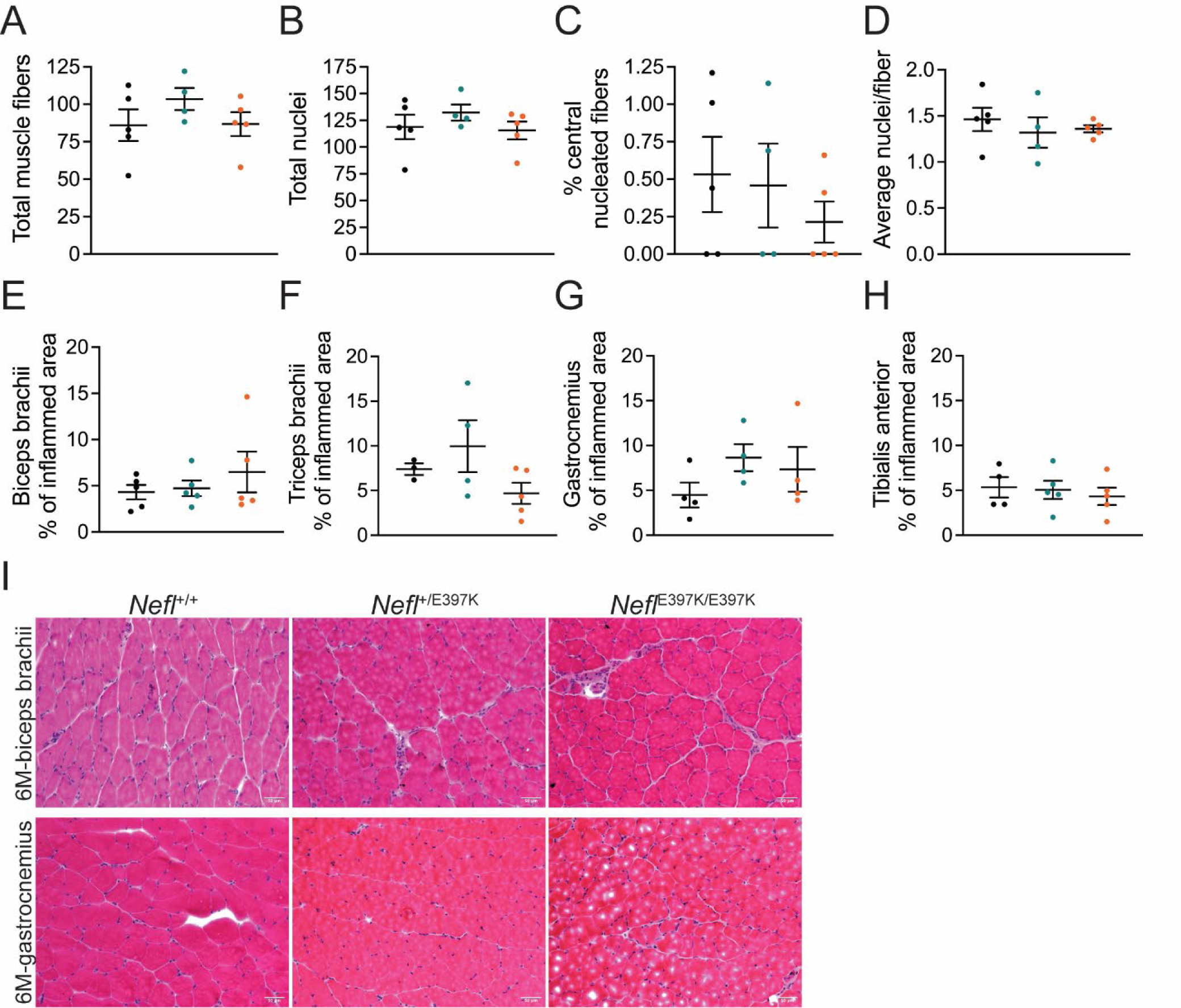
Hematoxylin and eosin (H&E) staining of CMT2E mice. Cross sections of biceps brachii, triceps brachii, gastrocnemius, and tibialis anterior muscles of *Nefl^+/+^* (black), *Nefl^+/E397K^* (teal), and *Nefl^E397K/E397K^* (orange) were stained with H&E. (A) Total muscle fibers in the gastrocnemius muscle at six months (*Nefl^+/+^*=86.0, *Nefl^+/E397K^*=102.5, *Nefl^E397K/E397K^*=86.8). (B) Total muscle fiber nuclei in the gastrocnemius muscle at six months (*Nefl^+/+^*=118.8, *Nefl^+/E397K^*=132.8, *Nefl^E397K/E397K^*=115.5). (C) Percentage of central muscle fibers in the gastrocnemius muscle at six months (*Nefl^+/+^*=0.53, *Nefl^+/E397K^*=0.54, *Nefl^E397K/E397K^*=0.21). (D) Average nuclei per muscle fiber in the gastrocnemius muscle at six months (*Nefl^+/+^*=1.5, *Nefl^+/E397K^*=1.3, *Nefl^E397K/E397K^*=1.4). (E) Six months biceps brachii percentage of inflamed area of *Nefl^+/+^* (4.3%, N=5), *Nefl^+/E397K^* (4.7%, N=5), and *Nefl^E397K/E397K^* (6.5%, N=5). (F) Six months triceps brachii percentage of inflamed area of *Nefl^+/+^* (7.5%, N=3), *Nefl^+/E397K^* (10.7%, N=4), and *Nefl^E397K/E397K^* (4.7%, N=5). (G) Six months gastrocnemius percentage of inflamed area of *Nefl^+/+^* (4.5%, N=4), *Nefl^+/E397K^* (8.7%, N=4), and *Nefl^E397K/E397K^* (7.4%, N=4). (H) Six months tibialis anterior percentage of inflamed area of *Nefl^+/+^* (5.7%, N=4), *Nefl^+/E397K^* (5.1%, N=5), and *Nefl^E397K/E397K^* (4.3%, N=5). (I) Representative images of the H&E staining at six months (6M) of biceps brachii (top panel) and gastrocnemius (bottom panel) muscles. Two-way ANOVA with Dunnett multiple comparison test was used for statistical analyses. There was not significance between any of the analyzed parameters. Data points/dots represent mice. N=number of mice evaluated.

## Notes

### Competing Interest Statement

Ownership: CLL is co-founder and chief scientific officer of Shift Pharmaceuticals. MAL is associated with Shift by family relation. Income: CLL has received more than $10,000 in income per annum from Shift Pharmaceuticals. Research support: Research in the CLL and MAL labs have been supported by sub-awards from Shift Pharmaceuticals (as part of grants from the DOD, CMT Research Foundation, and the NIH). Intellectual property: an invention disclosure has been submitted covering the intellectual property related to this work.

